# Differential sensitivity to LINE 1-induced damage contributes to the expansion of Tet2-deficient HSCs upon chronic inflammatory stress

**DOI:** 10.1101/2025.07.21.665900

**Authors:** Anne Stolz, Cheng Xuan, Christelle Freitas, Rabie Chelbi, Hassan Ait-Ougouram, David Fandrei, Mengliang Ye, Elisabeth Nelson, Claire Maillard, Mathieu Bohm, Salomeh El-Choufani, Weiwei Zhang, Audrey Porquet, Kahina Issoulaiene, Christian Muchardt, Elina Zueva, Michele Goodhardt, David Garrick, Françoise Porteu, Emilie Elvira-Matelot

## Abstract

Chronic inflammation disrupts hematopoietic stem cell (HSC) function and drives the expansion of TET2-mutated clones, fostering clonal hematopoiesis of indeterminate potential (CHIP). However, the molecular mechanisms linking inflammation to these hematopoietic alterations remain poorly understood. Here, we uncover a pivotal role of transposable element (TE) derepression, specifically of the most recent LINE-1 (L1) elements, in mediating inflammation-induced HSC dysfunction. We show that chronic exposure to low-dose lipopolysaccharide (LPS) causes a loss of the heterochromatin mark H3K9me3 at L1s in wild-type (WT) HSCs, leading to L1 activation, DNA damage, and reduced clonogenic capacity. Remarkably, *Tet2^−/-^* HSCs are resistant to this LPS-induced L1 derepression and associated genomic instability. Furthermore, targeted degradation of L1 RNAs in WT HSCs diminishes the competitive advantage of *Tet2^−/-^* HSCs under chronic inflammatory conditions both in vitro and in vivo. These findings reveal epigenetic control of L1 elements as a previously unrecognized mechanism that links chronic inflammation to HSC impairment and clonal expansion of *Tet2*-mutant cells. This work highlights TE regulation as a critical determinant of hematopoietic fitness and clonal evolution under inflammatory stress.

## INTRODUCTION

As cells age, they acquire somatic mutations due to environmental exposure and replication errors. Hematopoietic stem cells (HSCs), that give rise to all blood cells throughout life, are not spared by this phenomenon. While most of these mutations will have no functional consequence, some may lead to a clonal advantage and subsequent expansion. In the hematopoietic tissue, this phenomenon is referred to as clonal hematopoiesis of indeterminate potential (CHIP). CHIP is defined as the presence of somatic mutations with a variant allele frequency ≥2% in the peripheral blood of individuals without evidence of hematologic disease (Busque et al., 2012; Jaiswal et al., 2014; Genovese et al., 2014).

Most CHIP mutations are found in genes commonly mutated in myelodysplastic syndrome (MDS) and acute myeloid leukemia (AML), with mutations in epigenetic factors such as TET2, DNMT3A and ASXL1 being the most frequent (Jaiswal et al., 2014; Genovese et al., 2014; Kar et al., 2022; Bick et al., 2020). CHIP is associated with an increased risk of atherosclerosis and cardiovascular diseases (Fuster et al., 2017; Sano et al., 2018b; a) and with a 10-fold increased risk of leukemia (Jaiswal et al., 2014; Genovese et al., 2014; Bowman et al., 2018; Mitchell et al., 2021). It is thus considered as a preleukemic state. Around 75% of AML patients have a CHIP mutation years before diagnosis (Desai et al., 2018). Understanding the mechanisms by which CHIP mutations confer a selective advantage to HSCs and drive the emergence of preleukemic clones is therefore of critical importance.

The frequency of allelic variants at a given time depends both on the age at which the mutation appears (and the time the clone has had to expand) as well as on the clone’s ability to gain an advantage over non-mutated clones. The prevalence of CHIP increases thus with age and is present in more than 20% of people over the age of 70 (Jakobsen et al., 2024). However, longitudinal studies have demonstrated that the expansion of identical mutations varies among individuals (Young et al., 2016; Watson et al., 2020; Fabre et al., 2022; Pershad et al., 2025) and that environmental factors influence CHIP expansion. Several studies have investigated the mechanisms by which specific mutant clones gain an advantage over non-mutated HSCs and have led to a better understanding of the selective environments that drive their expansion (Florez et al., 2022). For example, clones with mutations in DNA repair pathways are selected and expand preferentially in response to genotoxic stresses, such as ionizing radiations, in the context of space exploration (Brojakowska et al., 2022; Mencia-Trinchant et al., 2020), or in radiotherapy and chemotherapy (Takahashi et al., 2017; Coombs et al., 2017; Bolton et al., 2020; Gillis et al., 2017; Desai and Roboz, 2019). In contrast, expansion of clones mutated in epigenetic factors, such as TET2, appears to be favored by the emergence of an inflammatory microenvironment in the bone marrow which is one of the characteristic features of aging (inflammaging) (Cook et al., 2020; Cai et al., 2018; Caiado et al., 2023; McClatchy et al., 2023). Further, since TET2 mutant HSCs display a myeloid-biased potential (Quivoron et al., 2011; Li et al., 2011; Moran-Crusio et al., 2011; Ko et al., 2011; Jakobsen et al., 2024) and give rise to differentiated blood cells with enhanced inflammatory properties (Yeaton et al., 2022; Pronier et al., 2022; Zhang et al., 2015; Sano et al., 2018b; Fuster et al., 2017), the expansion of these mutant HSCs is also likely to further promote the inflammatory microenvironment of the bone marrow in a feed-forward manner. This proinflammatory role has been linked to the higher risk of developing cardiovascular or autoimmune diseases (Cobo et al., 2022). *TET2* mutated HSCs are resistant to inflammatory stimuli and cytokines that can be produced by their myeloid progeny (Cai et al., 2018; Caiado et al., 2023; Abegunde et al., 2018; Jakobsen et al., 2024), and thus present a selective advantage which eventually leads to their expansion (Cook et al., 2020; Avagyan et al., 2021; Speck, 2022; Jakobsen et al., 2024). However, although one study has shown that under chronic IL-1β exposure, Tet2-deficient HSCs resist a remodeling of DNA methylation pattern observed in wild-type (WT) HSCs (McClatchy et al., 2023), the molecular mechanisms underlying the resistance of *TET2*-mutated HSCs in an inflammatory context still remain poorly understood.

Maintenance of HSC identity and differentiation potential is dependent on the trimethylation of lysine 9 of histone H3 (H3K9me3) (Koide et al., 2016; Djeghloul et al., 2016). One very important role of heterochromatin, through DNA methylation and H3K9me3, is the repression of transposable elements (TEs). TEs, which are further classified as either long terminal repeat (LTR) sequences (including endogenous retroviruses (ERV)), or non-LTR elements (including long and short interspersed elements (LINE-1/L1; SINE)), represent almost half of the human and mouse genomes. Originally thought of as junk DNA, they are gaining increasing interest, especially in aging and cancer which are strongly associated with chromatin disorganization. TEs constitute an important source of genomic instability since they can spread in the genome through a copy/paste mechanism and induce DNA damage (Gasior et al., 2006; Belancio et al., 2010; Barbieri et al., 2018). TEs also serve as a source of endogenous double-stranded (ds) RNA or cytoplasmic cDNA that are sensed as genomic parasites by members of the RIG-I-like (RLR) receptors or cGAS/STING pathways, eventually triggering the production of interferon type-I (IFN) and other inflammatory cytokines through activation of IRF3/NF-κB pathways (Gazquez-Gutierrez et al., 2021).

In HSCs, erosion of heterochromatin upon aging and stresses is accompanied by overexpression of TEs (Djeghloul et al., 2016; Hong et al., 2023; Clapes et al., 2021; Barbieri et al., 2018; Pelinski et al., 2022). TE expression upon aging and chemotherapies leads to intrinsic sterile inflammation, loss of HSC clonogenicity (Hong et al., 2023), and HSC exit from quiescence (Clapes et al., 2021). We further showed that the decreased H3K9me3 observed in HSCs upon irradiation stress mainly occurs at the evolutionarily recent and active L1s and LTRs (L1Md and IAP respectively in mouse) (Pelinski et al., 2022). TE overexpression contributes to irradiation-induced long-term DNA damage in HSCs and loss of HSC function which can be reversed by reverse transcriptase inhibitors (RTIs) (Barbieri et al., 2018). These findings suggest that age- or stress-induced alterations in heterochromatin and TE derepression may represent a common, central molecular mechanism underlying changes in HSC function. However, it remains unknown whether chronic inflammation relies on the same mechanisms and to what extent they drive the expansion of *TET2*-mutated cells.

In this study, we demonstrate that chronic inflammation, mediated by repeated low doses of lipopolysaccharide (LPS), triggers heterochromatin erosion, derepression of L1 elements and L1-induced DNA damage in wild-type HSCs. In contrast, *Tet2^−/-^* HSCs resist LPS-induced L1 epigenetic derepression and associated DNA damage, ultimately leading to their higher competitiveness.

## RESULTS

### Chronic inflammation leads to decreased H3K9me3 heterochromatin mark in HSCs

In order to investigate the contribution of inflammation to changes in heterochromatin, we used a mouse model of chronic inflammation mediated by serial injections of low doses of LPS, which has been shown to lead to impaired HSC function (Esplin et al., 2011).

In these experiments, WT mice were injected with 6 µg LPS every 2 days for 4 weeks (**Figure 1A),** and the effect of this treatment on the repressive histone modification H3K9me3 in HSCs was analyzed by immunofluorescence. Interestingly, we observed a significant decrease in global levels of H3K9me3 after LPS treatment in Lin^−^ Sca^+^ Kit^+^ (LSK) CD34^−^ Flk2^−^ HSCs (**Figure 1B**). This change was not due to a decrease in the total level of histone H3 which is rather increased in the presence of LPS (**Figure 1C)**. The reduction in H3K9me3 induced by LPS was still detectable in HSCs isolated from mice 8 weeks after the end of the LPS treatment, indicating that LPS induces a stable epigenetic change **(Figure 1D**). This parallels the long-term damage to HSC function, which is also maintained after removal of the inflammatory stimulus (Esplin et al., 2011; Takizawa et al., 2017). To determine if decreased H3K9me3 in LPS-treated HSCs could be a general feature of chronic inflammation, we analyzed HSC heterochromatin in a second model of chronic inflammation, the Db/Db obese mice (Chua et al., 1996). These mice carry a mutation in the *LepR* gene encoding the Leptin receptor, leading to rapid weight gain associated with systemic inflammation and a myeloid-biased hematopoietic compartment (Nagareddy et al., 2014). Immunofluorescence staining revealed a significant decrease in global levels of H3K9me3 in HSCs of Db/Db mice relative to that of control (Db/+) littermates **(Figure 1E**). A similar, although less pronounced, reduction in H3K9me3 was also observed in Lin^−^ Sca^+^ Kit^+^ CD34^+^ Flk2^−^ short-term reconstituting stem cells (ST-HSCs) of Db/Db mice. These results support our findings in LPS-treated mice and strongly suggest that chronic inflammation alone is sufficient to induce an impairment of H3K9me3 heterochromatin in HSCs. To investigate this further, we generated mixed bone marrow chimeras by reconstitution of lethally irradiated mice with a 1:1 ratio of bone marrow cells from wild-type mice (WT) and from mice deleted for *Tlr4*, encoding the Toll-like receptor 4, the pattern-recognition receptor responsible for the detection of LPS stimulation in HSCs (Takizawa et al., 2017). The mice were treated with LPS as described above, and the effect on H3K9me3 was analyzed by immunofluorescence in WT and *Tlr4*^−/-^ HSCs, which could be distinguished using CD45.1/CD45.2 isotypic markers (**Figure 1F**). Strikingly, while a decrease in H3K9me3 was observed in CD45.1^+^ WT HSCs, no decrease of H3K9me3 was detected in CD45.2^+^ *Tlr4*^−/-^ HSCs from the same LPS-treated mice, indicating that the LPS-mediated decline in HSC heterochromatin occurs via a direct interaction with Tlr4 receptors expressed by the stem cells. Altogether, these findings indicate that chronic inflammation directly impacts the chromatin state of HSCs, leading to a stable reduction of H3K9me3.

**Figure 1.**
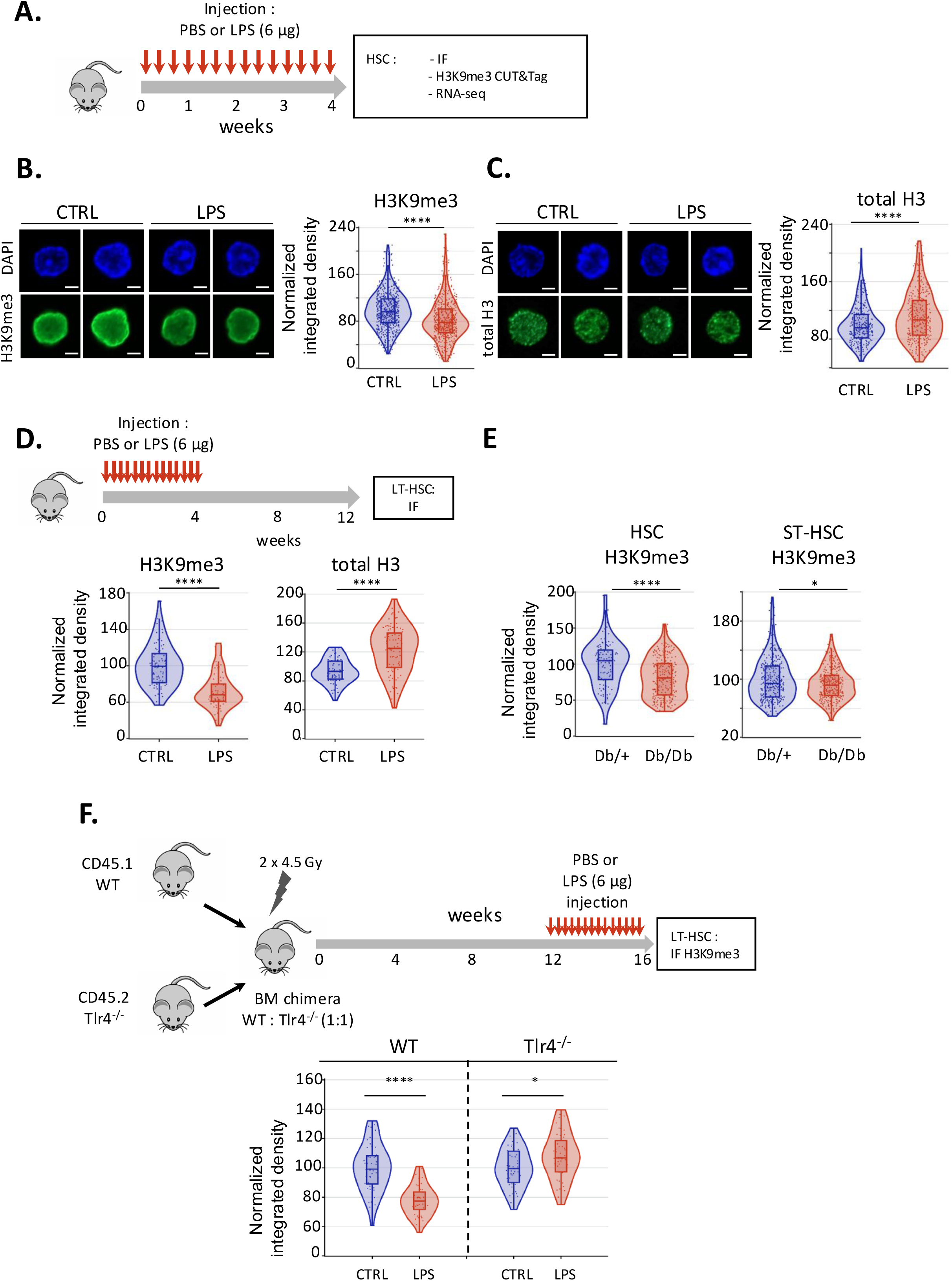
Chronic inflammation leads to decreased H3K9me3 heterochromatin in HSCs. **(A)** Schematic of the chronic inflammation model used in this study. Young mice were injected with either PBS (control-CTRL) or a low dose of LPS (6 µg) every 48 h over 4 weeks followed by immunofluorescence (IF), CUT&Tag or RNA-seq analysis of HSC. **(B-C)** Immunofluorescence analysis of H3K9me3 (B) and total histone H3 (C) in HSCs isolated from control or LPS-treated mice (minimum 9 animals per group). Representative images (left), and quantitation of fluorescent signal (right, minimum of 250 cells). **(D)** Mice were treated with PBS (control, n=3) or LPS (n=4) as in (A) and HSC were isolated at 8 weeks after completion of treatment for analysis of H3K9me3 (*left panel*) and total histone H3 (*right panel*) by IF. **(E)** IF analysis of H3K9me3 in HSC (left panel), and short-term (ST)-HSC (right panel) isolated from mice heterozygous (Db/+) or homozygous (Db/Db) for mutation of the *LepR* gene (3-6 animals per group). **(F)** (*Top*) Schematic of the experiment: Equal numbers of WT (CD45.1^+^) and Tlr4^−/-^ (CD45.2^+^) bone marrow cells were engrafted into lethally irradiated hosts. At 12 weeks post engraftment, mice were treated with PBS (CTRL, 3 mice) or LPS (4 mice) for 4 weeks as in (A), and HSCs were isolated for analysis of H3K9me3 by IF (*bottom*). For all quantitation of immunofluorescence, signal has been normalized to the mean value observed in the control group. Internal box plots indicate Q1, median and Q3 of each distribution. (**** p<0.0001; *** p<0.001; ** p<0.01; * p<0.05; ns indicates not significant; Mann Whitney).

### Chronic inflammation induces a loss of H3K9me3 at L1Md and IAP, the most recent subfamilies of L1s and LTRs

To further characterize the loss of H3K9me3 that occurs upon chronic inflammation, we performed H3K9me3 CUT&Tag experiments in HSCs sorted from LPS-treated mice (**Figure 1A).** We have previously shown that in HSCs, H3K9me3 is mainly enriched at TEs, and more particularly at L1Md elements, the most recent subfamilies of LINEs (Pelinski et al., 2022). In order to capture the information at these evolutionary youngest TE subfamilies, we performed a combined analysis of unique and multiple-mapping reads, as previously described (Pelinski et al., 2022). Focused differential enrichment analysis at TE genomic loci revealed 598 loci with significantly altered enrichment of H3K9me3, with 403 (67%) and 195 (33%) TEs showing decreased or increased H3K9me3 respectively upon LPS treatment (**Figure 2A and supplementary table S1).** Altogether across these 598 differentially altered TEs, there was a significant (p<0.0001) overall decrease in H3K9me3 concentration (**Figure 2B).** Among the TEs with decreased H3K9me3 following LPS treatment, there was a significant (p<0.0001-hypergeometric test) enrichment of LINEs (62.5%) and LTR (36.0%) as compared to the repartition of these TEs in the mouse genome (24.7% and 22.1% respectively) (**Figure 2C)**. LINEs were also significantly enriched in TEs gaining H3K9me3 (43.8%), but to a lesser extent than TEs which lose H3K9me3 (**Figure 2C)**. Consistent with these results, analysis across all 598 differentially affected TE loci confirmed a decrease in H3K9me3 at LINEs and at LTRs (**Figure 2D**). These data were confirmed in an independent H3K9me3 CUT&Tag experiment (**Figure S1 A-D** and **supplementary table S1**).

**Figure 2.**
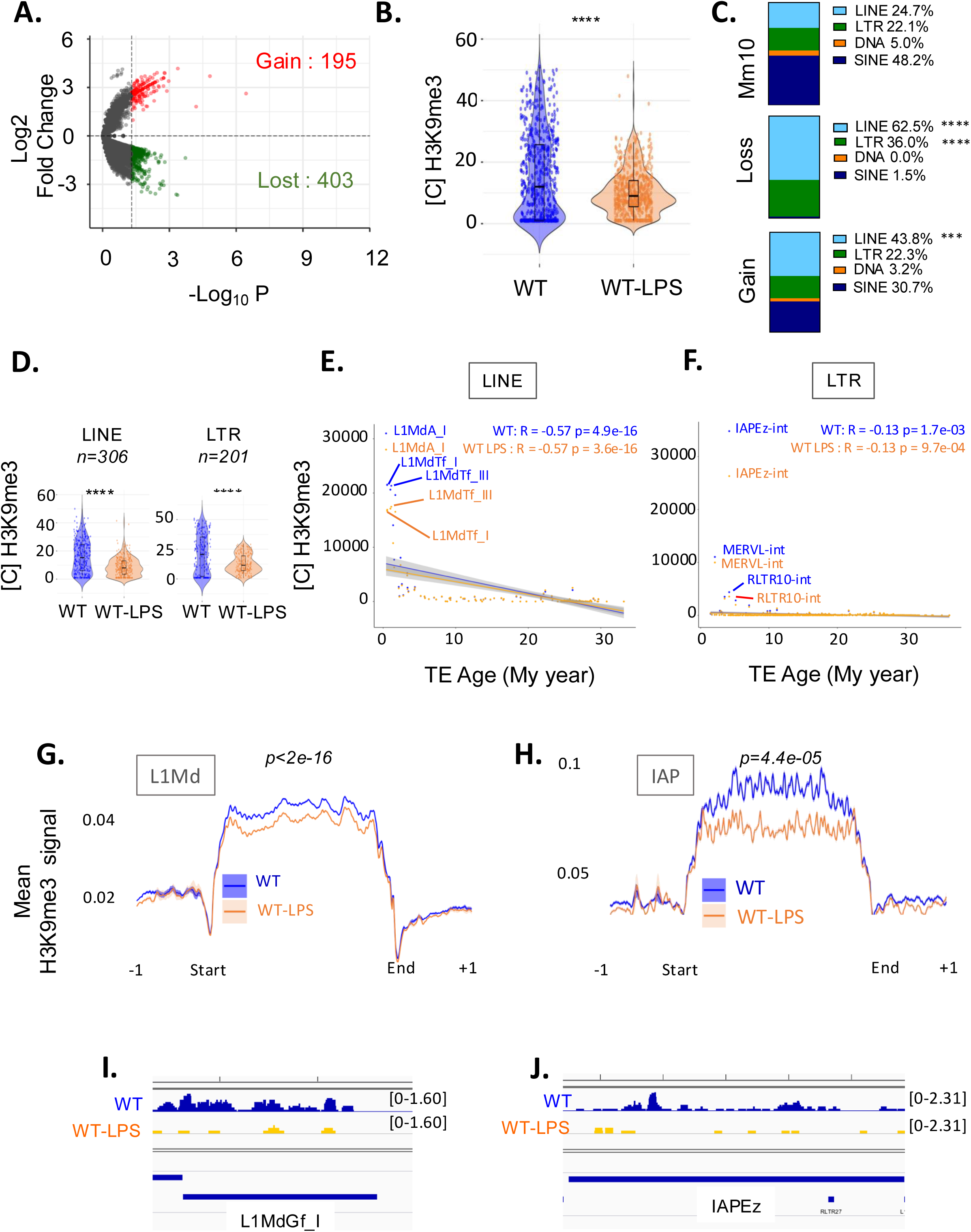
Chronic LPS treatment decreases H3K9me3 at the most recent subfamilies of LINEs and LTR in wild type HSC. **(A)** Volcano plot showing changes in H3K9me3 enrichment at TE loci in HSC between WT-LPS and WT conditions. TEs presenting a significant (*p<0.05)* increase (red) or decrease (green) of H3K9me3 enrichment which are analysed further in (B-D) are indicated. **(B)** Violin plot representing the distribution of H3K9me3 concentration at each TE locus retrieved in (A). **(C)** Repartition of the % of each family of TEs among the total TE loci in the mouse mm10 genome (*top*) and among the TEs retrieved in (A) presenting a significant decrease (*middle* - loss) or increase (*bottom* – gain) of H3K9me3. Hypergeometric test ***p<0.001, ****p<0.0001. **(D)** Violin plot representing the distribution of H3K9me3 concentration at each locus retrieved in (A) for LINE and LTR, SINE and DNA families of TEs in WT and WT-LPS conditions. Wilcoxon test. **p<0.01; ****p<0.0001. **(E-F)** Correlation plot representing H3K9me3 concentration quantified at all LINE (E) or LTR (F) elements *vs* their age in million years (My) in WT and WT-LPS conditions. R, Pearson correlation coefficient; p, pvalue. **(G-H)** Plot profile of the mean +/-SEM H3K9me3 enrichment along the L1Md (G) or IAP (H) sequences +/- 1kb flanking regions in WT-LPS (orange) *vs* WT (blue) conditions (2-3 samples per condition). **(I-J)** Integrative genomic viewer (IGV) visualization of H3K9me3 enrichment and TEs at one L1Md (L1Md_Gf chr8: 42202314-42203306) (G) and one IAP (IAPEz chr15: 29131255-29133920) (H) locus significantly losing H3K9me3 in WT (blue) vs WT-LPS (yellow) conditions. Merged of 2-3 samples per condition.

Since we and others have reported that evolutionarily recent TE subfamilies, particularly L1Md, are subject to tighter epigenetic regulation as compared to older TEs (Barau et al., 2016; Pezic et al., 2014; Pelinski et al., 2022), we further analyzed changes in H3K9me3 enrichment at LINEs and LTRs according to their evolutionary age, as calculated in (Sookdeo et al., 2013). As previously reported, we observed a negative correlation between the age of the LINE elements and H3K9me3 enrichment in WT HSCs, with the youngest repeats (typically L1Md_A, L1Md_Tf, and L1Md_Gf), showing the highest enrichment. Loss of H3K9me3 was mainly observed at these youngest subfamilies (**Figure 2E and S2A**). With respect to LTRs, while the anti-correlation between H3K9me3 enrichment and the age of LTRs is less pronounced, again the strongest enrichment of H3K9me3 was also observed at the youngest elements, especially IAPEz and MERVL (**Figure 2F and S2B**) and H3K9me3 loss in response to LPS treatment is mainly observed at these youngest subfamilies.

Finally, plot profiles of H3K9me3 enrichment across all L1Md and IAP sequences confirmed significant decrease of H3K9me3 at these elements in response to LPS treatment (**Figure 2G-H)** as exemplified on an integrative genome viewer **(Figure 2I-J)**. This decrease was particularly evident at elements which show the highest enrichment of H3K9me3 in the basal state: L1Md_A (**Figure S2C)** and IAPEz **(Figure S2D)** subfamilies. As previously described in embryonic stem cells (Bulut-Karslioglu et al., 2014), intact L1Md and IAP (conservation score > 97%), present a stronger H3K9me3 enrichment than degenerate elements (conservation score < 97%), and are the most affected by H3K9me3 loss upon LPS treatment (**Figure S2E-H).** Finally, H3K9me3 loss was specific for long L1Md (>5kb), that have more chance to contain the sequences encoding for the reverse transcription machinery (endonuclease and reverse transcriptase), compared to truncated L1Md (<5kb) **(Figure S2I-J).** Altogether, these data suggest that chronic LPS treatment induces a decrease of H3K9me3 at evolutionarily recent and intact subfamilies of L1s and LTRs that are the most likely to be damaging to the cell.

### Chronic inflammation induces a gain of H3K9me3 at L1Md in *Tet2^−/-^* HSCs

Since inflammation favors the expansion of *TET2*-mutated HSCs, we were prompted to compare these observations in WT HSCs with the effect of chronic inflammation on the heterochromatin compartment in *Tet2^−/-^*HSCs. Following application of the LPS injection protocol to *Tet2^−/-^*mice, H3K9me3 CUT&Tag experiments on sorted HSCs revealed 241 TE loci with significantly differential enrichment of H3K9me3 in response to LPS treatment. Surprisingly, by contrast with what was observed in WT HSCs, more loci (133, 55%) were observed to gain H3K9me3 following LPS treatment than those that lost it (108, 45%) (**Figure 3A and supplementary table S1).** Accordingly, the distribution of the H3K9me3 concentration at these differential peaks showed a slight but significant increase following LPS treatment (**Figure 3B).** Compared to the genomic distribution of TEs, there was a significant (p<0.001) enrichment of LTR (32.4%) among the TEs with decreased H3K9me3, while there was a significant (p<0.01) enrichment of LINEs (38.2%) among the TEs gaining H3K9me3 (**Figure 3C**). Consistent with this, analysis of H3K9me3 concentration at the 241 loci with differential H3K9me3 enrichment between *Tet2^−/-^-*LPS and *Tet2^−/-^* conditions confirmed a significant increase in H3K9me3 at LINEs, but no significant changes at other TE classes (**Figure 3D**). Gain of H3K9me3 was mainly observed at L1Mds (**Figure 3E)**

**Figure 3.**
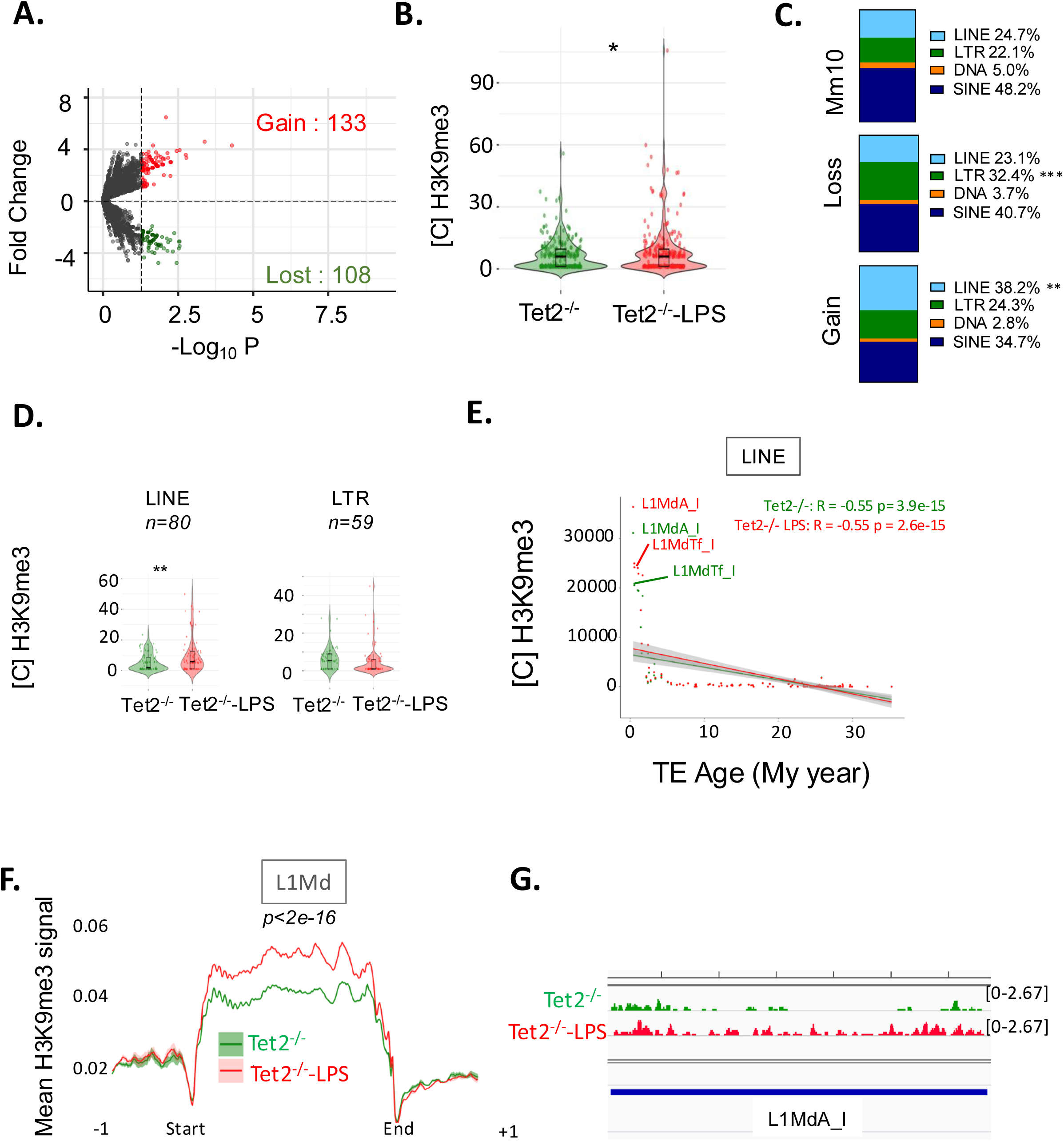
LPS induces a gain of H3K9me3 at L1Mds in *Tet2^−/-^* HSCs. **(A)** Volcano plot showing changes in H3K9me3 enrichment at TE loci in HSC between *Tet2^−/-^* mice +/- LPS. TEs presenting a significant (*p<0.05)* increase (red) or decrease (green) enrichment of H3K9me3 following LPS treatment (analysed further in (B-D)) are indicated. **(B)** Violin plot representing the distribution of H3K9me3 concentration at each TE locus retrieved in (A). **(C)** Repartition of the % of each family of TEs among the total TE loci in the mouse mm10 genome (*top*) and among the TE retrieved in (A) presenting a significant decrease (*middle* - loss) or increase (*bottom* – gain) of H3K9me3. Hypergeometric test **p<0.01; ***p<0.001 **(D)** Violin plots representing the distribution of H3K9me3 concentration at each locus retrieved in (A) for LINE and LTR families of TEs in *Tet2^−/-^*-LPS and *Tet2^−/-^* conditions. Wilcoxon test. **p<0.01; **(E)** Correlation plot representing H3K9me3 concentration quantified at all LINE elements *vs* their age in million years (My) in *Tet2^−/-^*-LPS *Tet2^−/-^*conditions. R, Pearson correlation coefficient; p, pvalue. **(F)** Plot profile of the mean +/- SEM H3K9me3 enrichment along L1Md sequences +/- 1kb flanking regions in *Tet2^−/-^*-LPS (red) *vs Tet2^−/-^* (green) conditions (2 samples per condition). **(G)** Integrative genomic viewer (IGV) visualization of H3K9me3 enrichment and TEs at one L1Md (L1MdA_I chr9: 40064126-40070688) locus significantly gaining H3K9me3 in *Tet2^−/-^*-LPS (red) vs *Tet2^−/-^* (green) conditions. Merge of 2 samples per conditions.

Finally, plot profile of H3K9me3 concentration along all L1Md sequences showed a significant increase of H3K9me3 at L1Mds upon LPS treatment in *Tet2*^−/-^ HSCs (**Figure F-G**). This increase was notably observed at intact rather than degenerate L1Mds (**Figure S3A-B)** and at both long (>5kb) and truncated (<5kb) L1Mds (**Figure S3C-D**). Altogether, these data show that LPS had inverse effects on H3K9me3 enrichment at L1 in WT and *Tet2*^−/-^ HSCs.

### Chronic LPS induces distinct transcriptomic changes in WT and *Tet2^−/-^* HSCs

To further investigate these different responses to LPS, we next compared the effect of chronic LPS treatment on the transcriptome in WT and *Tet2^−/-^* HSCs. Gene analysis revealed 332 differentially expressed genes (DEGs; p<0.05) upon LPS treatment in WT HSCs, of which 198 (60%) are down- and 134 (40%) are up-regulated **(Figure S4A)**, and 1260 DEGs found after LPS treatment of *Tet2^−/-^* HSCs, of which 746 (59%) are down-regulated and 514 (41%) are up-regulated (**Figure S4B**) (**supplementary table S2**). Of the 1541 genes whose expression was altered by LPS in either WT and *Tet2^−/-^*HSCs, only 51 genes (3.3%) were altered in both genotypes (**Figure S4C**), of which only 19 (1.2%) changed in the same direction (shown in green in **Figure S4D**).

We then performed Gene Set Enrichment Analysis (GSEA) on Hallmark gene sets (FDR<0.05, NES <-1.5 or >1.5) and showed that chronic LPS treatment led to inverse effects on gene pathways in WT and *Tet2^−/-^* HSCs. Indeed, while signatures of proliferative pathways (E2F and MYC targets) are downregulated following LPS treatment in WT HSCs (**Figure 4A-B),** they are positively enriched in *Tet2^−/-^* HSCs (**Figure 4A,C)**. We also observed inverse responses of the TNFA signaling via NFKB pathway upon LPS-treatment in WT (enriched) and *Tet2^−/-^*(depleted) HSCs (**Figure 4A,D-E)**.

**Figure 4.**
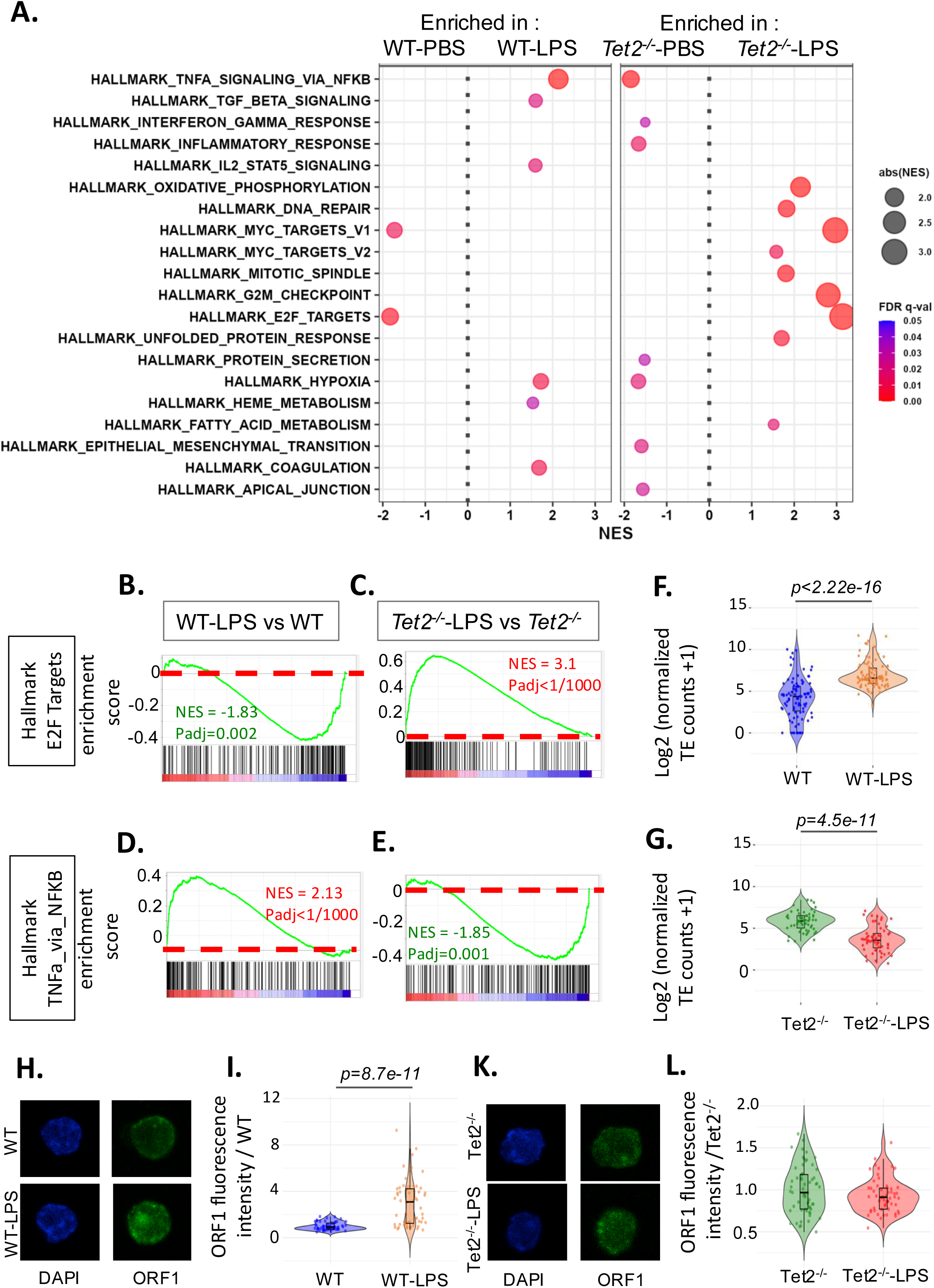
LPS induces differential transcriptomic responses in WT and *Tet2*^−/-^ HSCs. **(A)** GSEA analysis showing Hallmark gene sets significantly differentially enriched (padj<0.05 and -1.5<Normalized enrichment score (NES) or NES>1.5) in WT-LPS vs WT (left) and/or in *Tet2^− /-^*-LPS vs *Tet2^−/-^* (right) comparisons. **(B-E)** Enrichment plots for E2F TARGETS (B-C) and TNF-α signalling via NF-κB (D-E) hallmark genesets in WT-LPS vs WT (B,D) and *Tet2^−/-^*-LPS vs *Tet2^−/-^* (C,E) conditions. NES: normalized enrichment score. **(F-G)** violin plot representing TE subfamilies expression for each TE subfamily significantly (p<0.05) differentially expressed in WT-LPS vs WT (F) and *Tet2^−/-^*-LPS vs *Tet2^−/-^* (G) conditions. **(H-L)** Representative images (H, K) and quantification (I, L) of LINE 1 ORF1 fluorescent signal in HSCs sorted from WT (H,I) or *Tet2^−/-^* (K,L) mice after LPS treatment in vivo. Wilcoxon test. n=2 independent experiments, 1 mouse for each experiment, 25-33 cells measured per slide

Altogether, these data indicate that in comparison to WT HSCs, *Tet2^−/-^* HSCs show a distinct transcriptomic response to inflammation, associated with increased enrichment of proliferative pathways and decreased enrichment of inflammatory pathways.

We also analyzed the consequences of LPS-induced deregulation of H3K9me3 on TE expression in WT and *Tet2^−/-^* HSCs using TEtranscripts (Jin et al., 2015) (multiple mapping reads). LPS significantly (p<0.05) increased expression of 94 (97%) and decreased expression of 3 (3%) TE subfamilies in WT HSCs (**supplementary table S3**). In contrast, among the 57 significantly deregulated TE subfamilies upon LPS treatment in *Tet2^−/-^* HSCs, only 7 (12%) were up-regulated and 50 (88%) were downregulated (**supplementary table S3**). Altogether across the differentially expressed TEs, there was a significant (p<0.0001) overall increase in TE expression in WT (**Figure 4F)** but a significant decrease in *Tet2^−/-^* HSCs (**Figure 4G).**

When dealing with very recent TE subfamilies, it is difficult to precisely map reads originating from CUT&Tag and RNA-seq. We thus cannot compare our CUT&Tag and RNA-seq data to correlate H3K9me3 loss and TE expression at the level of individual loci. However, several L1 subfamilies that show inverse epigenetic changes in WT and *Tet2^−/-^* HSC upon LPS treatment, also showed differential expression, being significantly up- and down-regulated by LPS in WT and *Tet2^−/-^* HSC respectively (**Figure S4E,F**). Further, analysis at the protein level by IF revealed that in vivo LPS treatment specifically induced LINE ORF1 protein in WT but not in *Tet2^−/-^* HSCs (**Figure 4H-L).** Altogether, these data indicate that chronic LPS treatment increases the expression of active L1s specifically in WT but not in *Tet2^−/-^* HSCs.

### Derepression of TEs upon LPS treatment in WT HSC is associated with the accumulation of DNA damage and HSC functional changes

We and others have shown that L1 derepression and mobilization in HSCs are associated with double strand breaks (DSBs), as measured by the presence of γH2AX foci (Gasior et al., 2006; Belancio et al., 2010; Barbieri et al., 2018). Inflammation has also been shown to lead to genomic instability in HSPCs/HSCs (Takizawa et al., 2017; Rodriguez-Meira et al., 2023). L1s of greater than 5kb in length and bearing promoter sequences, could encode an intact ORF2 protein with endonuclease and reverse transcriptase activities which can potentially induce DNA damage. In order to characterize the consequences of the epigenetic derepression of these L1s in WT HSCs upon LPS treatment, we assessed γH2AX foci by immunofluorescence in HSCs sorted from mice at the end of chronic LPS treatment in vivo (**Figure 1A**), or in HSCs from untreated mice following culture in the presence or absence of 10µg/ml LPS for 48h in vitro. In both cases, LPS treatment induced a significant increase in the number of gH2AX foci in HSCs (**Figure 5A-D)**. In the in vitro conditions, the induction of gH2AX foci by LPS was inhibited in the presence of lamivudine (3TC), a reverse transcriptase inhibitor known to act on L1 reverse transcriptase activity (**Figure 5C-D**), suggesting that L1 is responsible for DNA damage induced by LPS in HSCs.

**Figure 5.**
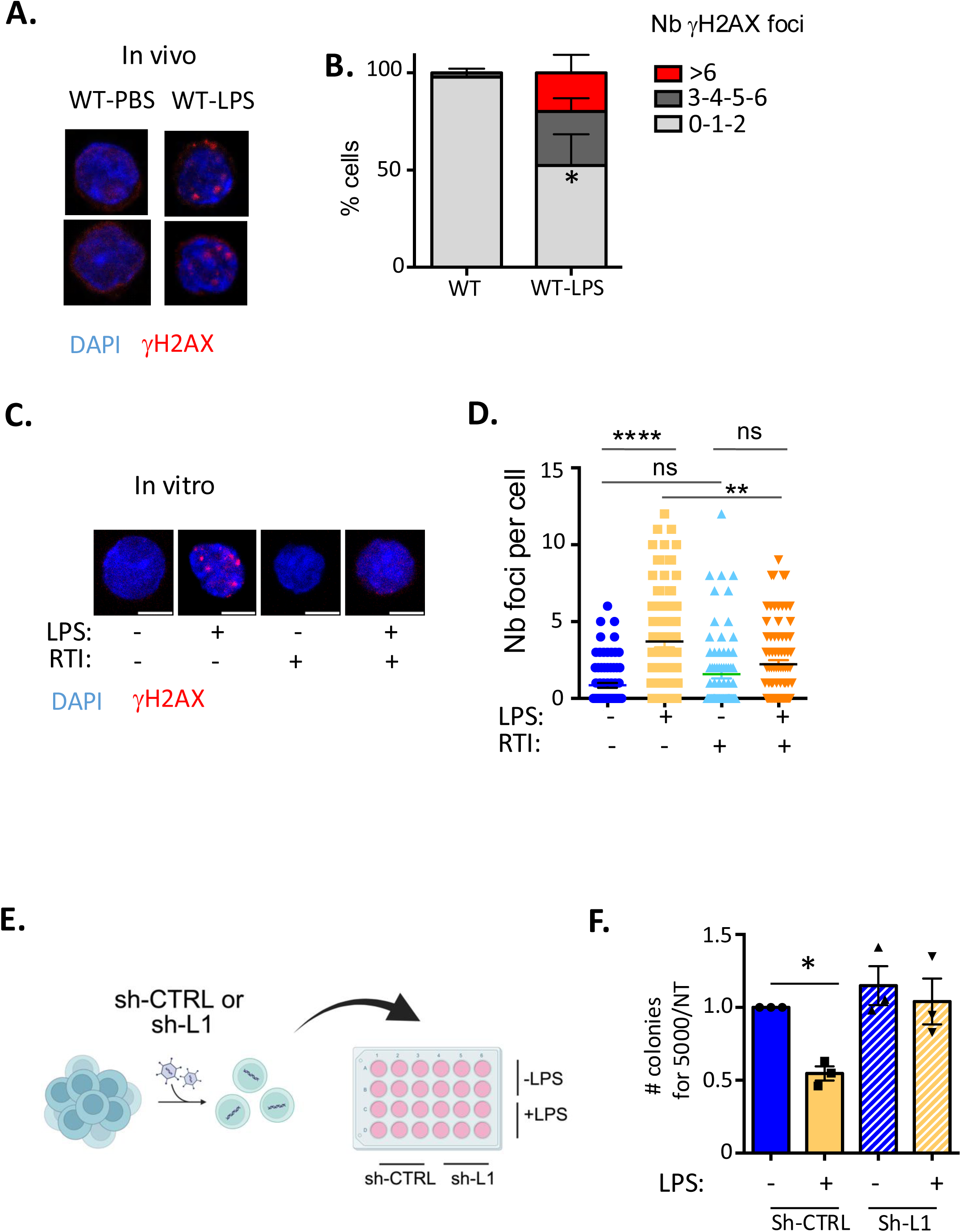
TE derepression upon LPS treatment in WT HSC is associated with the accumulation of DNA damage and loss of HSC function. **(A-B)** Representative images (A) and quantification (B) of gH2AX foci in HSCs sorted from WT mice treated or not with LPS in vivo. n=2 independent experiments, pools of 2 mice for each experiment, 38 to 55 cells counted per slide. 2 way ANOVA with Sidak’s multiple comparison test *p<0.05 for 0-1-2 foci per cell. **(C-D)** Representative images (C) and quantification (D) of gH2AX foci in HSCs +/-10µg/ml LPS and +/- 10µM lamivudine after 48h of culture in vitro. n=2 independent experiments, 3 pools of 2 mice for each experiment, 40-42 cells counted per slide. 1-way ANOVA with Tukey’s multiple comparison test. **** p<0.0001 **,p<0.01, ns: non-significant **(E)** Experimental design. Lineage – (Lin-) cells were transduced with either an sh-control or an sh-L1 and mcherry+ HSCs were sorted and plated in methylcellulose in the presence or absence of LPS 3 days after transduction. Figure created with biorender **(F)** Number of colonies formed from 5000 HSCs normalized to untreated cells. N=3 independent experiments. *p<0.05 1-way ANOVA with Tukey’s multiple comparison test.

To further assess whether L1-induced DNA damage affects HSC function upon LPS treatment, we transduced Lin^−^ cells with either an sh-control or an sh-L1 fused to mCherry reporter gene, that was previously shown to reduce L1Md expression in HSPCs (Clapes et al., 2021). We then plated mCherry positive HSCs in methylcellulose, in the presence or absence of 10µg/ml LPS, and assessed their clonogenicity (**Figure 5E).** While LPS induced a significant decrease in the number of colonies formed by HSCs transduced with an sh-control, it did not affect the clonogenicity of HSCs transduced with an sh-L1 (**Figure 5F).**

Altogether, these data suggest that loss of H3K9me3 at TEs, notably at L1s, contributes to the DNA damage and decrease in clonogenicity observed in HSCs in response to chronic inflammation.

### *Tet2^−/-^* HSCs are protected from the L1-mediated deleterious effects of LPS

We then assessed whether epigenetic changes observed at L1 elements in *Tet2^−/-^* HSCs may prevent the accumulation of DNA damage and functional alterations. Indeed, in contrast with findings in WT HSCs, LPS did not increase gH2AX foci in *Tet2^−/-^* HSCs both *in vitro* in culture and *in vivo* after chronic injections (**Figure 6A-C**). As reported previously (Esplin et al., 2011), treatment with LPS decreased the reconstitution capacity of WT HSCs (**Figure S5A-D**). To assess how TET2 loss affects the response of HSC to LPS in vivo, hematopoietic chimeras were created by transplanting CD45.1 recipient mice with 0.3 × 10^6^ bone marrow (BM) cells from CD45.2 WT or *Tet2^−/-^* mice (10%) in competition with 2.7 × 10^6^ BM (90%) cells from CD45.1 WT mice. At 1 month post-engraftment, recipient mice were treated with PBS (control) or LPS every two days for 30 days and reconstitution was assessed in BM and peripheral blood (PB) at the end of 5 months (**Figure 6D**). LPS treatment had no significant impact on peripheral blood engraftment of either WT or *Tet2^−/-^* cells after 5 months (**Figure S5E**). However, the proportion of WT CD45.2^+^ cells in LSK and HSC populations (**Figure 6 E-F**), as well as their absolute numbers (**Figure 6 G-H)** were reduced following LPS treatment as compared to PBS-treated mice. As previously described (Cai et al., 2018; Caiado et al., 2023), *Tet2^−/-^* HSCs present a significantly higher reconstitution capacity than WT HSCs (**Figure 6 E, Figure S5E**). In contrast to WT HSCs, the proportion of CD45.2^+^ *Tet2^−/-^* in LSK and HSC populations (**Figure 6 E-F**) as well as their absolute numbers (**Figure 6 G-H)** were not affected by LPS treatment. To determine the functional consequences of LPS exposure on long-term HSC function, 5.0 × 10^6^ BM cells from primary transplants were engrafted to secondary CD45.1 recipients. The effect of LPS on WT HSCs was still observed in these secondary transplantations, indicating a long-term influence on HSC function, while the reconstitution capacity of *Tet2^−/-^*HSCs was again unaffected (**Figure 6 I**).

**Figure 6.**
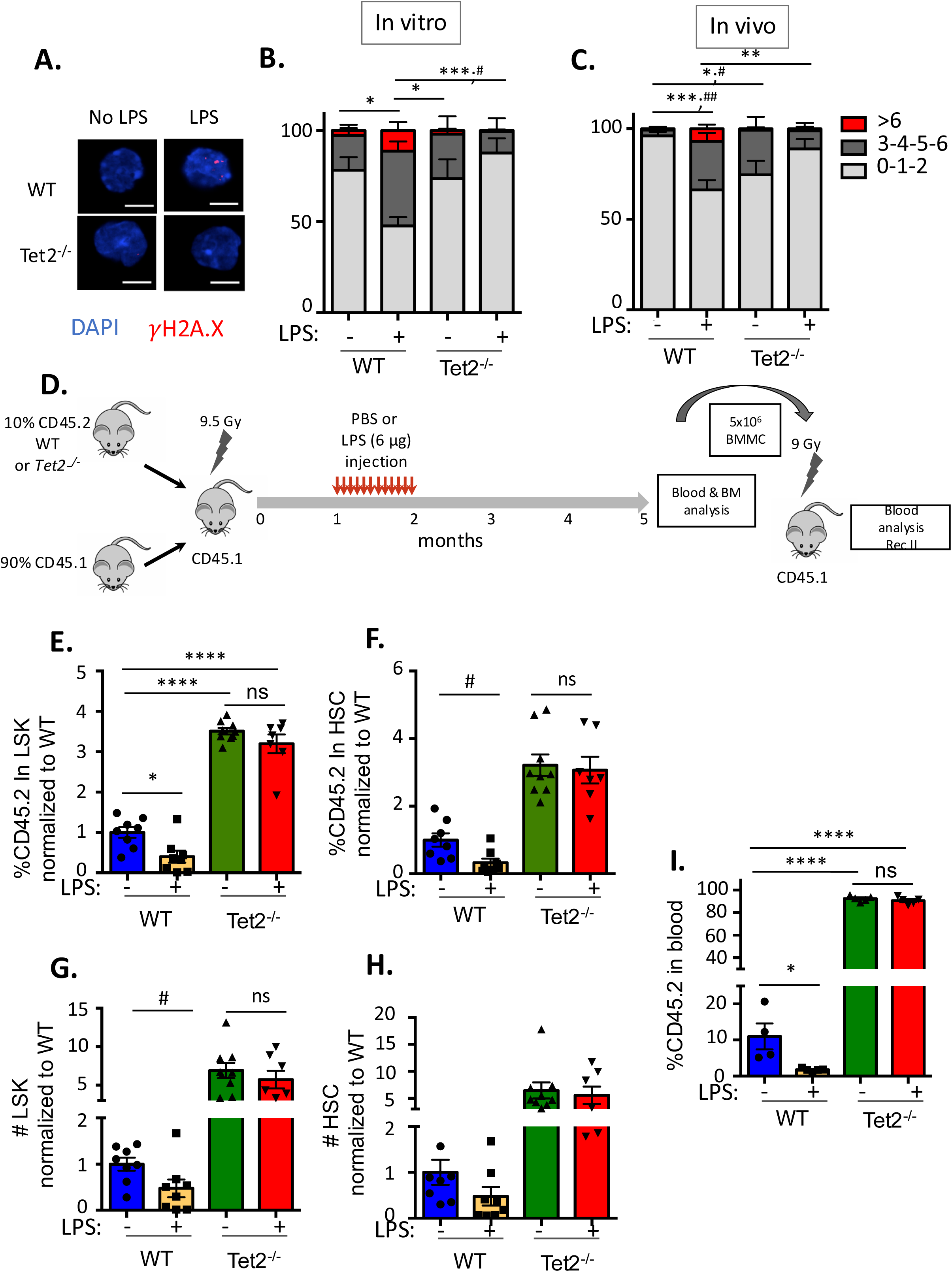
*Tet2*^−/-^ HSCs are resistant to the deleterious effects of LPS. **(A-C)** representative images (A) and quantification (B-C) of gH2AX foci in WT or *Tet2^−/-^* HSCs +/- 10µg/ml LPS in vitro (B) and after PBS or LPS treatment in vivo (C). n=3 independent experiments, 1 pool of 2 mice in each, 38 to 52 cells counted per slide. 2-way ANOVA with Tukey’s multiple comparisons test. *p<0.05; **p<0.01, ***p<0.001 for 0-1-2 foci comparisons; #p<0.05, ##p<0.01 for 3-4-5-6 foci comparison. **(D)** Experimental design for testing the response of WT or *Tet2^−/-^* cells to LPS treatment in chimeric mice in vivo. BMMC : Bone marrow mononuclear cells. **(E-F)** % CD45.2 in LSK (E) and HSCs (F) and **(G-H)** absolute numbers of LSK (G) and HSCs (H). **(I)** % CD45.2 in blood upon secondary reconstitution. One dot represents a single mouse. n=2 independent experiments *p<0.05; **p<0.01, ***p<0.001, ****p<0.0001 1-way ANOVA with Tukey’s multiple comparison test; #p<0.05 unpaired two-tailed t-test. ns = non significant.

### The competitive advantage of *Tet2^−/-^* HSCs upon LPS treatment is dependent on LINE1 expression in WT HSCs

The resistance of *Tet2^−/-^* HSCs to the deleterious effects induced by L1 activation may contribute to their clonal expansion following LPS treatment. To examine this hypothesis, we conducted in vitro competition assays between wild-type (WT) HSCs transduced with either a control shRNA (sh-control) or an L1-targeting shRNA (sh-L1) fused to an mCherry reporter, and *Tet2^−/-^* HSCs transduced with a GFP-labeled sh-control construct. We then monitored the relative expansion of *Tet2^−/-^* HSCs (GFP⁺) among the total transduced population (GFP⁺ and mCherry⁺ cells) after three days of culture in the presence or absence of LPS (**Figure 7A**). The results showed that *Tet2^−/-^*HSCs exhibited a significant expansion in the presence of LPS when competing with WT HSCs expressing the sh-control. However, this effect was abolished when WT HSCs expressed sh-L1 (**Figures 7B and S6A**).

**Figure 7.**
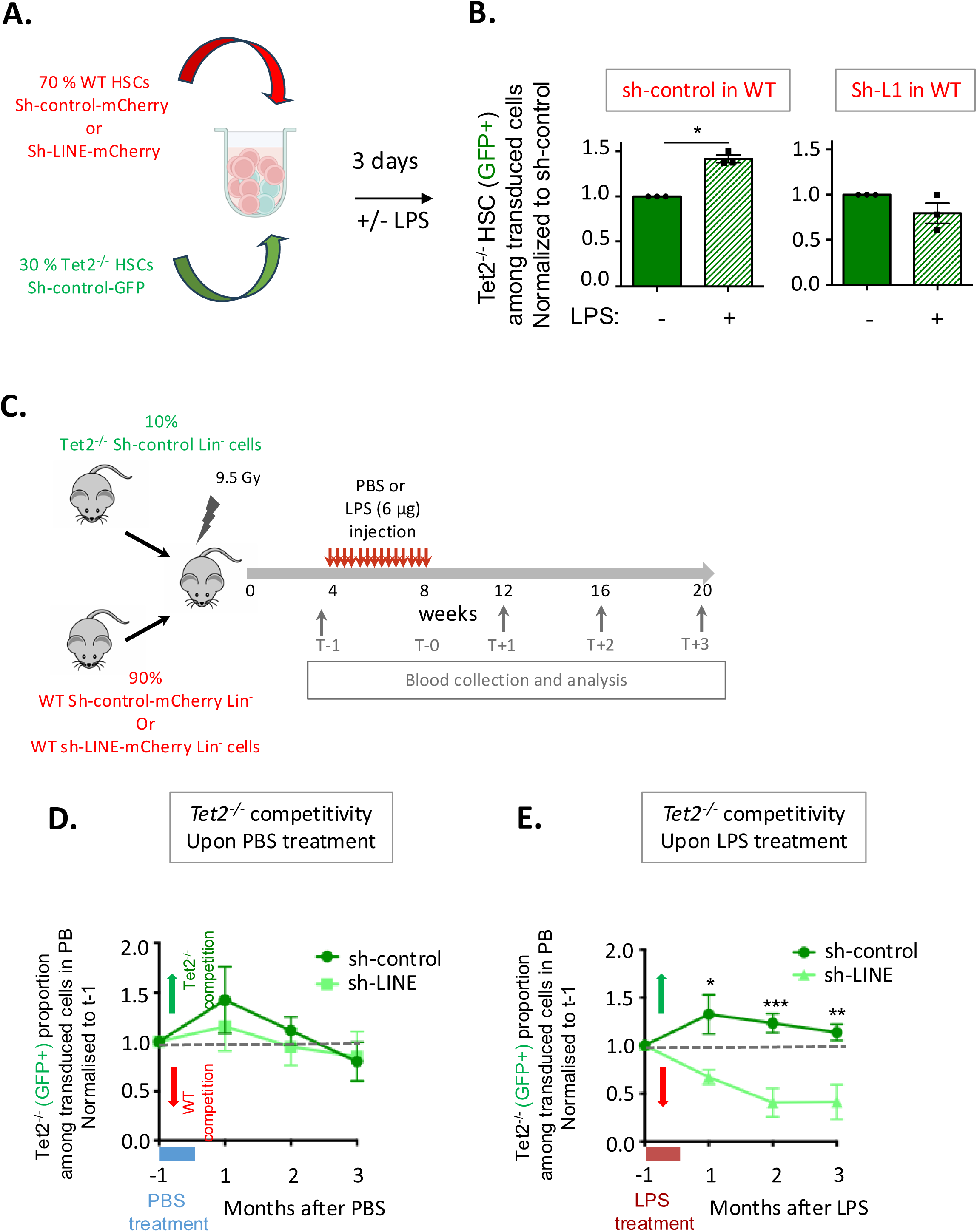
The competitive advantage of *Tet2^−/-^* HSCs upon LPS treatment is dependent on LINE1 expression in WT HSCs. **(A)** Experimental design to analyze *Tet2^−/-^* vs WT HSCs competition in vitro in the presence or absence of LPS. 70% of WT HSCs transduced with either an sh-control or sh-LINE fused to mCherry reporter gene were cultured in vitro with 30% of *Tet2^−/-^* HSCs transduced with an sh-control fused to GFP reporter gene, in the presence or absence of LPS. **(B)** Proportion of *Tet2^−/-^ HSCs* (GFP+) among the total number of cells (GFP+ or mCherry+) when in competition with WT HSCs transduced with a sh-control (left) of with a sh-L1 (right) and in the presence (+) or absence (-) of LPS in the culture medium. Results are normalised to the non-LPS condition. *p<0.05 paired t-test. **(C)** Experimental design to analyze *Tet2^−/-^* vs WT HSCs competition in vivo. 10% of CD45.2 lin-cells transduced with an sh-control-GFP were engrafted with 90% of CD45.2 lin-cells transduced with an sh-L1-mcherry or an sh-control-mcherry in CD45.2 recipient mice previously irradiated with 9.5 Gy. Recipient mice received chronic PBS or LPS injections during 30 days. Blood collections were performed before and monthly after treatment. **(D-E)** proportion of *Tet2^−/-^* cells (GFP+) in total transduced cell subsets (GFP+ and mCherry+) when in competition with WT cells transduced with sh-control (dark green) or with sh-L1 (light green in peripheral blood at 1 to 3 months after PBS (D) or LPS (E) treatment, normalized to the proportion of *Tet2^−/-^* cells measured before starting the treatment. Above or below the dotted line means that *Tet2^−/-^* cells are more or less competitive than WT respectively. N=2 independent experiments. 5-7 mice per condition. Multiple unpaired t-test. *p<0.05, **p<0.01, ***p<0.001

To validate these findings in vivo, we transplanted a mixture of hematopoietic progenitor cells composed of 10% Tet2⁻/⁻ Lin⁻ cells transduced with a GFP-labeled sh-control and 90% WT Lin⁻ cells transduced with either an mCherry-labeled sh-control or sh-L1 construct into lethally irradiated CD45.2 recipient mice. One month post-transplantation—defined as time point T-1—mice received either LPS or PBS injections (**Figure 7C**). In each animal, we quantified the proportion of *Tet2^−/-^* cells (GFP⁺) among total engrafted cells (GFP⁺ and mCherry⁺) in peripheral blood (PB) both before (T-1) and monthly after LPS or PBS treatment (T1 to T3). The GFP⁺ cell proportion at each time point was normalized to the pre-treatment (T-1) level (**Figures 7C and S6B**).

In PBS-treated mice, no difference was observed in the proportion of *Tet2^−/-^* cells when competing with WT cells expressing either sh-control or sh-L1 (**Figures 7D and S6B**). In contrast, LPS-treated mice showed a significant reduction in the competitive advantage of *Tet2^−/-^* cells when the competing WT population expressed sh-L1 compared to sh-control (**Figures 7E and S6B**).

Together, these results indicate that the clonal expansion of *Tet2^−/-^* HSCs under chronic inflammatory conditions depends on the presence of L1 transcripts and their deleterious impact on WT HSCs (**Figure 8**).

**Figure 8.**
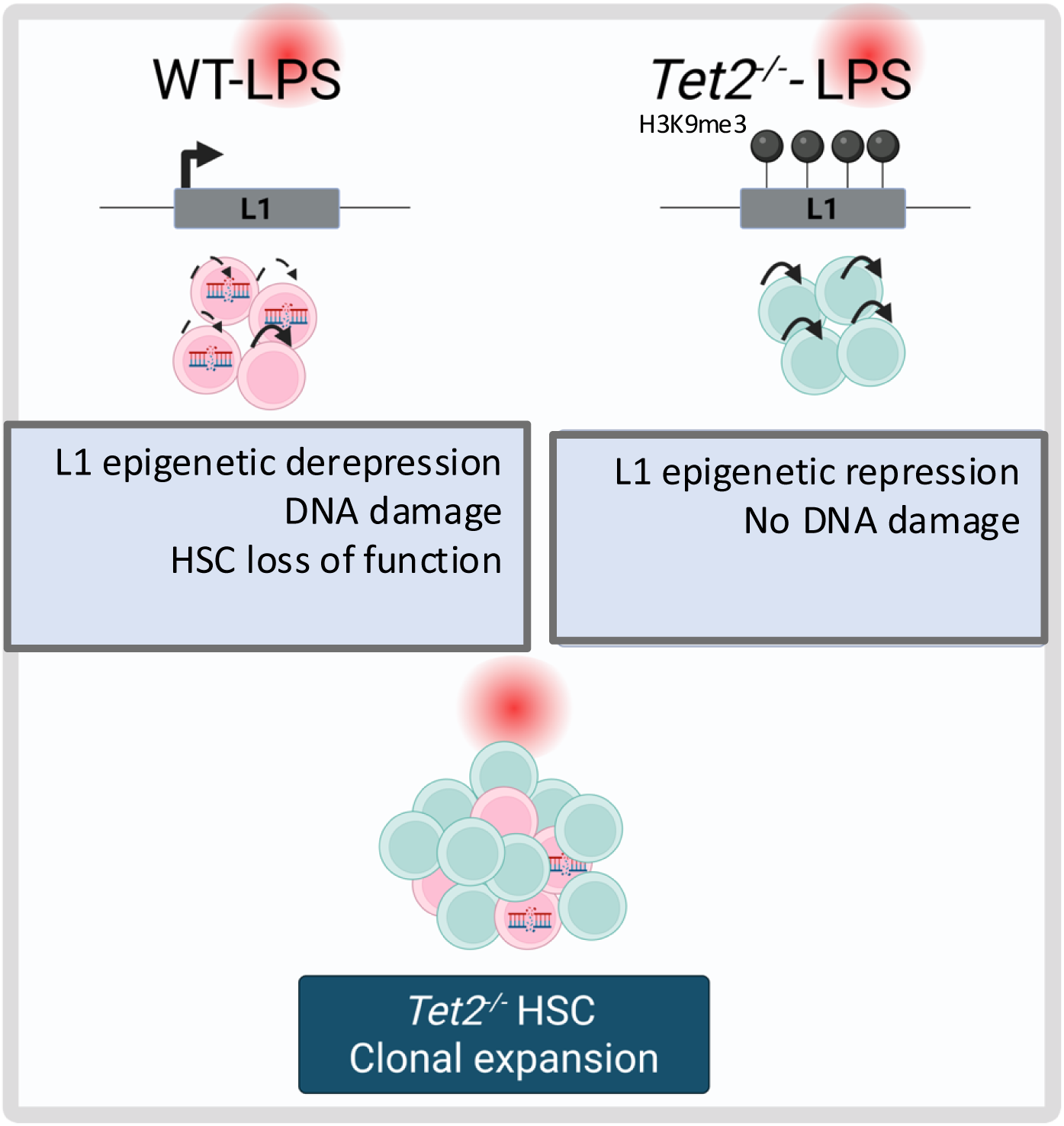
Model. Upon chronic inflammatory stress, the function of WT HSCs is altered through loss of H3K9me3 at L1, L1 overexpression and L1-induced accumulation of DNA damage. *Tet2^−/-^*are resistant to L1-induced deleterious effects upon LPS treatment through epigenetic repression, eventually leading to their clonal expansion. Figure created with biorender.

## DISCUSSION

Through their ability to modify the transcriptome and to induce genomic instability, TEs can exert an oncogenic activity (Chuong et al., 2017; Liang et al., 2024). However, when expressed above a certain threshold, TEs can induce DNA damage, senescence, and a state of “sterile inflammation” and thus play the role of tumor suppressors (Zhao et al., 2021; De Cecco et al., 2019). We previously showed that derepression of young active L1Md, which occurs upon H3K9me3 loss, is partly responsible for the sustained DNA damage which is observed in HSCs even a long-time after irradiation. Preventing this damage with RTIs is associated with a significant improvement of HSC reconstitution capacities (Barbieri et al., 2018; Pelinski et al., 2022). Our present findings add to previous studies showing the accumulation of DNA damage and genomic instability upon inflammation in HSCs/HSPCs (Takizawa et al., 2017; Rodriguez-Meira et al., 2023). We further show here that L1 epigenetic derepression following H3K9me3 loss contributes to DNA damage in HSCs and their decreased clonogenicity upon chronic inflammation. We also show that *Tet2^−/-^* HSCs are refractory to LPS-induced epigenetic derepression of L1 elements and L1-induced DNA damage, giving them an advantage compared to WT HSCs, and eventually leading to their clonal expansion under inflammatory conditions.

Global heterochromatin loss, and more particularly redistribution of heterochromatin, was proposed as a theory of aging (Tsurumi and Li, 2012). We previously reported that the H3K9me3 heterochromatin mark is depleted in HSCs of aged mice and humans. This is correlated with increased expression of TEs such as L1Mds and IAPs (Djeghloul et al., 2016). Similarly, Hong et al reported a relocalization of heterochromatin in mouse HSPCs, correlated with increased expression of LTR elements (Hong et al., 2023). However, HSC aging is a complex and multifactorial process in which chronic inflammation represents only one of multiple contributing features. Our current study shows that chronic inflammation alone is sufficient to induce a decrease of H3K9me3 in HSCs and the epigenetic derepression of TEs. We further show that chronic inflammation leads to a relocalization of H3K9me3 at the different TE classes in HSCs, with a decreased concentration at some LINEs and LTR but an increased enrichment at some SINEs and DNA transposons. Interestingly it has been shown that compared to HSCs, myeloid progenitors exhibit a relocalization of H3K9me3 from the periphery to the center of the nucleus (Ugarte et al., 2015). As inflammation pushes HSCs towards myeloid differentiation, it would be of interest to study the role played by H3K9me3 relocalization and TE derepression on the induction of a myeloid program in HSCs.

Selective pressures that arise during aging modulate the function of HSC and eventually favor the emergence of competitive clones capable of self-renewal in the aged bone marrow. These emerging clones also display a predisposition to oncogenic transformation. We show here that the differential epigenetic regulation of L1 in *Tet2^−/-^* vs WT HSCs and the resistance of *Tet2^−/-^*HSCs towards TE-induced DNA damage may contribute to the clonal expansion of *Tet2^−/-^* HSCs observed under inflammatory stress. Our findings add to previous studies showing that *Tet2^−/-^*HSCs are resistant towards epigenetic changes such as heterochromatin relocalization and loss of DNA methylation, which are observed upon aging and chronic IL1β treatment, respectively (Hong et al., 2023; McClatchy et al., 2023). In our study of a more restricted HSC population, we did not observe a differential regulation of LTRs in *Tet2^−/-^* and WT mice, as previously observed in HSPCs (Hong et al., 2023). We rather observed a decrease in H3K9me3 enrichment at LTRs in both WT and *Tet2^−/-^* HSCs upon LPS treatment. However, we observed that LPS induced opposing effects at intact L1s, which displayed decreased and increased H3K9me3 enrichment in WT and *Tet2^−/-^* HSCs respectively. Using sh-L1, we further demonstrate that L1s are necessary for the decreased competitivity of WT HSCs relative to *Tet2^−/-^* HSCs.

It has previously been shown in mouse embryonic stem cells that members of the HUSH complex are required for specific repression of L1 elements, and particularly the most recent L1Mds, by recruiting the SETDB1 methyltransferase and H3K9me3 (Robbez-Masson et al., 2018; Liu et al., 2018). Inactivation of members of the HUSH complex leads to impaired proliferation of an AML cell line through L1-induced DNA damage (Gu et al., 2021). However, our RNA-seq analysis did not reveal a significant upregulation of members of the HUSH complex, such as Tasor, Mphosph8 (mpp8) or Pphln1 (periphilin 1), that could account for the specific increase in H3K9me3 enrichment at L1s in *Tet2^−/-^* HSCs upon LPS treatment (**supplementary table2**).

Previous studies have shown that the survival advantage of TET2-deficient HSCs under inflammatory conditions is linked to changes in anti-vs pro-apoptotic gene expression, increased expression of DNA repair pathways, and an increase in proliferative and HSC self-renewal transcriptomic signatures (Cai et al., 2018; Abegunde et al., 2018; Caiado et al., 2023; McClatchy et al., 2023). As compared to WT HSCs, *Tet2^−/-^* HSCs also show resistance to DNA demethylation of transcription factor binding sites involved in differentiation, which could contribute to their enhanced self-renewal capacity (McClatchy et al., 2023). We did not observe a maintenance of HSC signatures in *Tet2^−/-^* HSCs upon LPS treatment as compared to WT in our RNA-seq data (data not shown). However, we confirm that LPS induced an increase in DNA repair pathways in *Tet2^−/-^* HSCs, which was not observed in WT cells (**Figure 4A**). In addition to the epigenetic repression of L1, this upregulation of DNA repair genes may thus also play a role in the resolution of DNA damage accumulation in *Tet2^−/-^*HSCs and their resistance towards LPS treatment. Finally, as previously observed in TET2-mutated vs WT HSCs in individuals with CHIP (Jakobsen et al., 2024), our data show an attenuated transcriptional response of *Tet2^−/-^* HSCs towards inflammation, with a significant decrease of the TNFA_signalling_via_NFKB pathway. Hence, in addition to epigenetic repression of L1, this attenuated response may further increase the clonal advantage of *Tet2^−/-^* HSCs, as previously shown in Tet2 mutant HSC in zebrafish (Avagyan et al., 2021). Interestingly, we also observed a significant decrease in interferon gamma response pathway in *Tet2^−/-^* HSCs (**Figure 4A**), that could be the consequence of TE repression.

Following from our study, it will be of interest to investigate whether repression of TEs contributes to the predisposition to oncogenic transformation which is characteristic of Tet2-mutant HSCs. Interestingly, expression of TEs, and notably L1s, are repressed in leukemic stem cells from patients with AML (Colombo et al., 2017; Gu et al., 2021). Further, in patients with chronic myelomonocytic leukemia (CMML), 60% of whom present with biallelic mutations of TET2, we recently showed an increase of the repressive H3K9me2 mark at TEs and a repression of immune associated transcripts. This could allow the resistance and further expansion of mutated cells as compared to their WT counterparts upon aging (Hidaoui et al., 2024). Epigenetic silencing of TEs by the HUSH complex and SETDB1 H3K9 methyltransferase is required for AML cell line proliferation, by preventing TE-induced DNA damage and interferon response (Gu et al., 2021; Cuellar et al., 2017). Our current findings suggest that TE repression could be an important early event contributing to clonal evolution and the establishment of a pre-leukemic state.

Altogether, the findings of this study have revealed a novel epigenetic mechanism that favours clonal expansion of *Tet2^−/-^* HSC under conditions of inflammatory stress and which may contribute to further oncogenic evolution of expanded clones.

## METHODS

### Mouse strains and treatment

*Tet2^−/-^* and WT littermates (2-4 months age) were provided by Olivier Bernard (INSERM U1170 – Gustave roussy) (Quivoron et al., 2011). Db/Db mice (BKS(D)-Leprdb/db/JOrlRj) and Db/+ controls (BKS(D)-Leprdb/+/JOrlRj) (14 weeks old) were obtained from Janvier Lab. TLR4^−/-^ mice were provided by Christian Muchardt (Institut de Biologie Paris Seine). C57BL/6 CD45.2 or CD45.1 mice (6-8 weeks old) were from Envigo or Charles River laboratories respectively. All the mice were housed in a specific pathogen-free environment. All procedures were reviewed and approved by the Animal Care Committee N°26 (APAFIS #23286-2019121214392327 and APAFIS #35147-2022020315419972). *Tet2^−/-^* and WT littermates received intraperitoneal injections of 6 µg LPS-B5 Ultrapure (Invivogen, tlrl-pb5lps) every 2 days for 28 days and control mice were injected with 1x PBS (adaptation of the protocol described by Esplin et al. (2011)). Mice were irradiated with 9.5 Gy with an X-ray irradiator (RX irradiator X-RAD 320) or with a cesium irradiator (GSR D1^®^) for reconstitution experiments with total BM, or shRNA-transduced Lin-cells, respectively. Engraftments were performed by retro-orbital injections.

### Cell harvest and culture

BM was harvested from femur, tibia and hip bones. Total BM was depleted of differentiated hematopoietic cells (lineage-positive; lin+ cells) using Mouse Hematopoietic Progenitor Stem Cell Enrichment Set (BD Biosciences) and LT-HSCs (Lin-, Sca+, c-Kit+,CD34-, Flk2-) or ST-HSCs (Lin-, Sca+, c-Kit+,CD34+, Flk2-) were sorted using ARIA3, ARIA Fusion or Influx cell sorters (BD Franklin Lakes, NJ, USA), using the following antibodies: CD3ε (Lin) – APC clone 145-2C11 (553066, BD), TER-119 (Lin) – APC clone Ter-119 (557909), CD45R/B220 (Lin) – APC clone RA3-6B2 (553092), Ly6G-6C (Lin)-APC clone RB6-8C5 (553129), Ly-6A/E (Sca-1) - PeCy7 clone d7 (558162), CD117 (c-Kit) – PE or PerCP-Cy5.5, clone 2B8 (553355 or 560557 respectively), CD34– FITC or AF700 clone RAM34 (560238 or 560518), CD135 (Flk2) – BV421 or PE clone A2F10.1 (562898 or 553842 respectively), all from BD Biosciences, and collected in StemSpan (StemCell Technologies).

Freshly isolated murine HSCs were cultured in StemSpan medium supplemented with 1% penicillin/streptomycin (P/S), containing the following cytokines (all from Miltenyi Biotech): FTL3-Ligand (FTL3-L, 10ng/ml), stem cell factor (SCF, 100ng/mL) interleukin 3 (IL3, 10ng/mL) and interleukin 6 (IL6, 10ng/mL), at 37°C, 5% CO2. For treatments, 10µg/mL LPS-B5 Ultrapure (Invivogen, tlrl-pb5lps) or 10µM lamivudine (Sigma, 3TC-L1295-10MG) were added at the start of the cultures (day 0) for 48h.

### Immunofluorescence

For H3K9me3 analysis: sorted HSCs were pipetted onto poly-lysine-coated slides (2.000-5.000 cells per spot), and allowed to fix to the slide for 1 h at room temperature in a humid chamber. Cells were then fixed with 2% formaldehyde / PBS for 20 min and permeabilized in 0.5% Triton X-100 / PBS for 15 min at room temperature. After washing three times with PBS, cells were blocked for 1 h in 5% BSA / 5% goat serum / 0.05% Tween / PBS and were incubated with anti-H3K9me3 (07-523, Millipore) or anti-H3 (ab18521, Abcam) primary antibodies diluted in blocking solution overnight at 4°C in a humid chamber. Slides were washed three times in PBS and incubated for 1 h at room temperature in 1% BSA / PBS containing anti-rabbit-Alexa488 (A11008, Invitrogen). After three washings in PBS, slides were mounted using VectaShield containing DAPI (H1200, Vector Laboratories). Stained cells were analyzed using a 63× objective lens Zeiss Lsm 800 Confocal microscope and Zeiss Zen blue software. Mean fluorescence intensity (Integrated density) of primary antibodies was measured using ImageJ software.

For analysis of DNA damage and ORF1 signal, 3000-5000 HSCs were cytospun onto glass slides and immunofluorescence was performed as previously described (de Laval et al., 2013b). γH2AX antibody was purchased from Millipore (05-636-I) and used at 1/2000, ORF1 antibody was purchased from Abcam (EPR21844-108) and used at 1/1000. Detection was performed using Alexa Fluor 555 anti-mouse (Invitrogen, A-21425) or Alexa Fluor 488 anti rabbit (Invitrogen, A11008) secondary antibody (1/600). Slides were visualized using an SPE or SP8 confocal microscope (Leica).

### Cell transduction with sh-L1

Lentiviral vectors expressing control sh-RNA (scramble sequence: ATCTCGCTTGGGCGAGAGTAAGTAGTGAAGCCACAGATGTACTTACTCTCGCCCAAGCGAGAG) or shRNA targeting L1 (tgctgttgacagtgagcgCTAAACAGGTGATGGAAATGATAGTGAAGCCACAGATGTATCATTTCCATCACCTGTTTAGtgcctactgcctcgga), together with GFP or mCherry reporter genes, were kindly provided by Dr. Antonio Morales-Hernandez (University of Michigan). Lentivirus was prepared using a three-plasmid system: (1) shRNA transfer vector-, (2) Gag/Pol-+ Rev/Tat-(pCMV), and (3) envelope plasmid (pMD2.G) by cotransfection of HEK293T cells using jetPRIME® (OZYME, POL101000046). Supernatant was collected 48 h later, cleared, titred onto HEK293T cells and stored at −80 °C. 50µL of virus were used to transduce 2 million Lin-cells and incubated for 72h before sorting of GFP+ or mCherry+ HSCs.

### CFU assays

350 HSCs (Lin-, Sca+, c-Kit+,CD34-, Flk2-) transduced with sh-L1 or sh-control were treated or not with 10µg/mL LPS-B5 Ultrapure (Invivogen, tlrl-pb5lps)LPS for 48h in liquid culture before seeding in 24-well plates containing 500µl of methylcellulose (M3434, StemCell), supplemented with 1% P/S (ThermoFischer Scientific). Colonies were counted after 7 days.

### CUT&Tag

CUT&Tag-IT assay kit (Active Motif) was used on 3,000 HSCs according to manufacturer’s instructions. Cells were incubated O/N with 0.5µg of H3K9me3 (C15410193-Diagenode). Transposed DNA fragments were amplified 18-fold by PCR using adapters supplied with the kit. PCR purification was carried out using the same Kit to remove remaining primers and large fragments. The quantity and quality of the libraries were assessed on Agilent 2100 Bioanalyzer (Agilent Technologies 50567-4626). Sequencing of the libraries was performed on the NovaSeq-6000 at Gustave Roussy (Illumina; 50 bp paired-end reads).

### CUT&Tag Analysis

Quality control was performed using FastQC (v0.11.9) and MultiQC (v1.11). Reads were aligned to the mm10 (UCSC genome) reference using Bowtie2 (v2.4.1) with the following parameters: --end-to-end --very-sensitive --no-mixed --no-discordant --phred33 -I 10 -X 700. Duplicate reads were removed with Picard (v2.26.9) using the parameters --REMOVE_DUPLICATES true --VALIDATION_STRINGENCY LENIENT. In classical pipelines, multiple mapping reads are usually discarded by applying a MapQ quality score threshold of 30 (MapQ>30) to aligned reads. In our pipeline, in order to retain multiple-mapping reads, we have eliminated this threshold and applied MapQ>0 using Samtools (v1.13). By definition, this approach will retain both unique and multiple mapping reads. Aligned reads were sorted and indexed with Samtools (v1.13). A coverage track (bigWig) was generated using deeptools (v3.5.0) with parameters -bs 5 --normalizeUsing BPM. All bam and bigwig files were generated by downsampling of each sample to that of the sample with the lowest number of reads

The Bioconductor package Diffbind (Ross-Innes et al., 2012) (v3.8.4) was employed in R v4.2.3 to quantify H3K9me3 levels at TEs, using a reconstructed version of the RepeatMasker annotation which allows the assembly of different fragments of an element (Bailly-Bechet et al., 2014; Walter et al., 2016). Paired-end mode was enabled during the read counting step using the SummarizeOverlaps method. The default mapping quality threshold (mapQCth) was adjusted to 0 for multimapping analysis. Differential binding affinity was assessed using the DBA_DESEQ2 method. The normalized H3K9me3 concentration across all TE loci within the same family or subfamily was aggregated to calculate the total H3K9me3 concentration per TE family. Differential binding at peaks was determined using a P-value threshold of 0.05. Correlation plots of TE age and H3K9me3 concentration was plotted with ggplot2 (v3.5.1) in R (v4.2.3), assuming a genomic substitution rate of 1.1% per million years (Sookdeo et al., 2013) and summing the mean H3K9me3 concentration at TEs from the same subfamily. A coverage track (bigWig) was generated using deeptools’ bamCoverage with the following parameters: -bs 5 -normalizeUsing BPM. Scores over different genomic regions were calculated using deeptools’ computeMatrix with the following parameters --beforeRegionStartLength 1000 --afterRegionStartLength 1000 --regionBodyLength 2500 missingDataAsZero. The mean density profile for 2-3 samples was then plotted with the SEM using the R package collection tidyerse (Wickham et al., 2019). Statistics of the signal were calculated using the R package Rseb (Gregoricchio et al., 2022) and the Wilcoxon test.

### RNA-seq

HSCs from individual mice were lysed in Tri-Reagent (Zymo Research) and stored at −80°C until used. Total RNA was extracted using the Direct-Zol RNA microprep kit (Zymo research). All samples were subjected to SMARTer® Stranded Total RNA-Seq Kit v3 - Pico Input Mammalian (Takara Bio USA, Inc., 634485) following the manufacturer’s instructions. The fragmentation time was adjusted according to RNA quality, after evaluation on the Agilent Bioanalyzer. Libraries were pooled and sequenced (2 × 100bp) on the NovaSeq6000 (Illumina).

### RNA-seq analysis

Quality control was performed using FastQC (v0.11.9) and MultiQC (v1.11). UMI was extracted using umi_tools (v1.1.2). Adapters were removed with Trimgalor (v0.6.5). The FASTQ file was filtered for rRNA sequences using SortMeRNA (v3.4.6). Reads were aligned to the GRCm38 mouse genome reference using STAR (v2.7.5a) and gene quantification was performed using salmon (v1.6.0). Differential expression analysis was performed using DESeq2 R package with P-value <0.05.

For pathway enrichment analysis and data visualization, we performed Gene Set Enrichment Analysis (GSEA) using the GSEA command line (v 4.3.3, Broad Institute) (Subramanian et al., 2005). Normalized counts from the DESeq2 differential expression analysis were used as input. Enrichment analysis was performed against the mm10 Hallmark gene sets obtained from the Molecular Signatures Database (MSigDB) (Castanza et al., 2023). Default parameters were used, except for the permutation type which was set to “gene_set” and metric set to Diff_of_Classes.

Enrichment scores and statistical significance were calculated using 1000 gene set permutations. Results were considered significant at a false discovery rate (FDR) of less than 0.05. GSEA results were plotted and visualized using the R package tidyverse.

### TETranscripts

To allow sufficient number of multireads during alignment for TE analysis, --outFilterMultimapNmax and --winAnchorMultimapNmax were set to 100. TE quantification was performed using TETranscripts (v2.2.3) with the parameter mode set to multi. Differential expression analysis was performed using the DESeq2 R package with P-value <0.05.

CUT&Tag and RNA-seq analysis were performed on the Core Cluster of the Institut Français de Bioinformatique (IFB) (ANR-11-INBS-0013).

### Statistical analysis

Results were statistically evaluated using GraphPad Prism version 9.0 software (GraphPad Software Inc.). The results are displayed as the means and SEM. The value of *, P < 0.05 was considered as significant, and **, P < 0.01 or ***, P < 0.001 as highly significant.

## Non-standard abbreviations

HSC: hematopoietic stem cell
CHIP: clonal hematopoiesis of indeterminate potential
MDS: myelodysplastic syndromes
AML: acute myeloid leukemia
CMML: chronic myelomonocytic leukemia
TE: transposable element
LTR: long terminal repeat
ERV: endogenous retrovirus
LINE/SINE: long/short interspersed elements

## Data availability

The dataset generated from the CUT&Tag experiments for Figures 2, 3 and S1 are available in ArrayExpress accession E-MTAB-15445, E-MTAB-15444 and E-MTAB-15448 respectively and from the RNA-seq for Figures 4 in ArrayExpress accession E-MTAB-15446 and E-MTAB-15447.

## Acknowledgments

We thank animal facility, the Genomic and the Imaging and Cytometry Platforms of Gustave Roussy for CUT&Tag and RNA sequencing and cell sorting and confocal analysis, respectively. FACS analysis was supported by DIM-ITAC Région Ile de France. We also thank S. Duchez and members of the Platforme Technologique of the Institut de Recherche Saint Louis. We thank O. Bernard (INSERM U1170, Gustave Roussy) for providing *Tet2^−/-^* mice, C. Muchardt (Institut de biologie paris Seine) for providing *Tlr4^−/-^*mice, A. Morales-Hernandez (University of Michigan, US) for providing sh-L1, C. Marty (INSERM U1287, Gustave Roussy) for her advice on cell transduction and S. Lotersztajn for advice and fruitful discussion concerning the Db/Db mouse model. We also thank A. Teissandier and D. Bourc’his for providing us the reconstructed repeatMasker database. This work was supported by INSERM and grants from Ligue National Contre le Cancer (LNCC, Equipe labellisée EL2020) and ARC Foundation (n° ARCPGA2023110007352_7985) to F. P. and Agence National de la Recherche (ANR-15-CE14-0003 to M.G., ANR-23-CE14-0017 to F.P., and ANRJCJC20-CE14-0018-01 to E.E.M), and ARC Foundation (ARCPJA2022070005354), Fondation de France (n° 00135490/ PR-151961) and EHA kick off grant (EHA KOG 2022) to E.E.M.

A.S. is recipient of a fellowship from LNCC, C.X. is recipient of a fellowship from the Chinese Scientific Council, C.F. is supported by ANR (ANR-15-CE14-0003 to M.G. and C.M.) and A.P is a recipient from the Ministère de l’Enseignement Supérieur de la Recherche et de l’Innovation. M.B is supported by INCA (PLBIO N°2020-095 to F.P.) and ANR (ANR-23-CE14-0017 to F.P.), R.C is supported by ANR (ANRJCJC20-CE14-0018-01 to E.E.M) and ANR (ANR-23-CE14-0017 to F.P.), H.A.O is supported by ARC foundation (n° ARCPGA2023110007352_7985 to F. P.), and M.Y is supported by ANR (ANR-23-CE14-0017 to F.P.).

The authors declare no conflict of interest.

## Author contributions

A.S., C.X., C.F., W.Z., K.I, C. Ma and A.P. performed cytometry, IF experiments and analysed the results. A.S. and S.EC performed CUT&Tag and RNA-seq experiments. A.S., C.F., E.N. S.EC and E.E.M performed reconstitution experiments and analyzed the results. A.S. performed in vitro competition assays and analyzed the results. M.B. and C.X. performed CFU experiments and analyzed the results. R.C., D.F., H.A.O, M.Y, E.Z. and E.E.M. performed bioinformatic analyses. M.G. designed and supervised the initial studies in WT HSCs, and analyzed the results, D.G., F.P. and E.E.M. designed and supervised the study, analyzed the results and wrote the manuscript. M.G., C.M., F.P., and E.E.M. contributed to funding acquisition.

