## Supplementary material for "Differential sensitivity to LINE 1-induced damage contributes to the expansion of Tet2-deficient HSCs upon chronic inflammatory stress": Figures S1 to S6

FIGURE S1

Related to figure 2

Stolz et al.

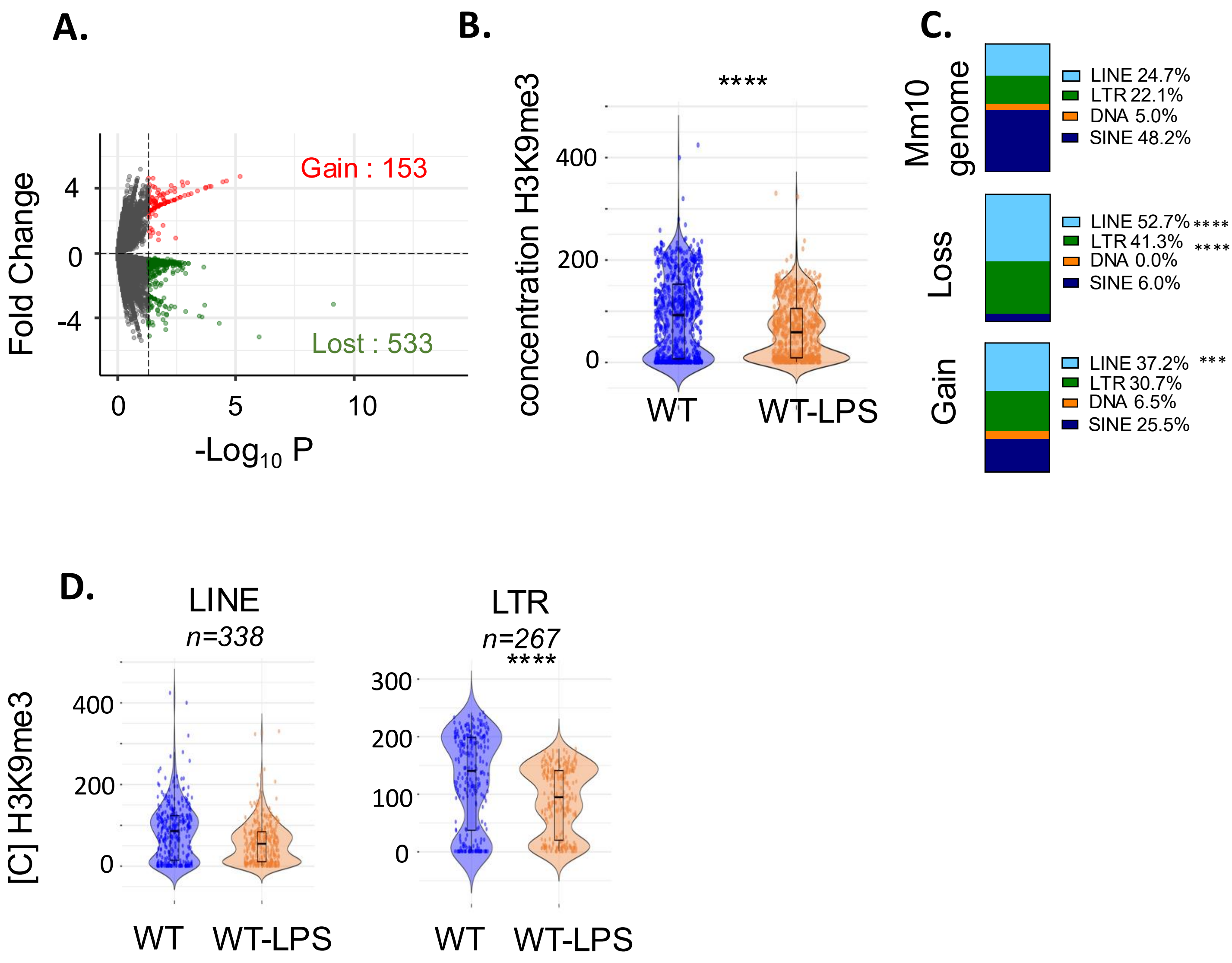


**Figure S1 (related to figure 2). (A-D)** Analysis of an independent H3K9me3 CUT&Tag experiment in HSC of WT-LPS and WT. **(A)** Volcano plot of the TE differentially enriched in H3K9me3 between WT-LPS and WT conditions. The horizontal axis represents the –log10(pValue) and the vertical axis the log2 fold change (FC). TE presenting a significant (*p<0.05)* increase (red) or decrease (green) enrichment in H3K9me3 are shown. **(B)** Violin plot representing the distribution of H3K9me3 concentration at each TE locus retrieved in (A). **(C)** Repartition of the % of each family of TE among the total TE loci in the mouse mm10 genome (*top*) and among the TE retrieved in (A) presenting a significant decrease (*middle* - loss) or increase (*bottom* – gain) of H3K9me3. Hypergeometric test. ***p<0.01, ****p<0.0001 **(D)** Violin plot representing the distribution of H3K9me3 concentration at each locus retrieved in (A) for LINE and LTR families of TEs in WT and WT-LPS conditions. Wilcoxon test. ****p<0.0001.

FIGURE S2-Related to figure 2 Stolz et al.

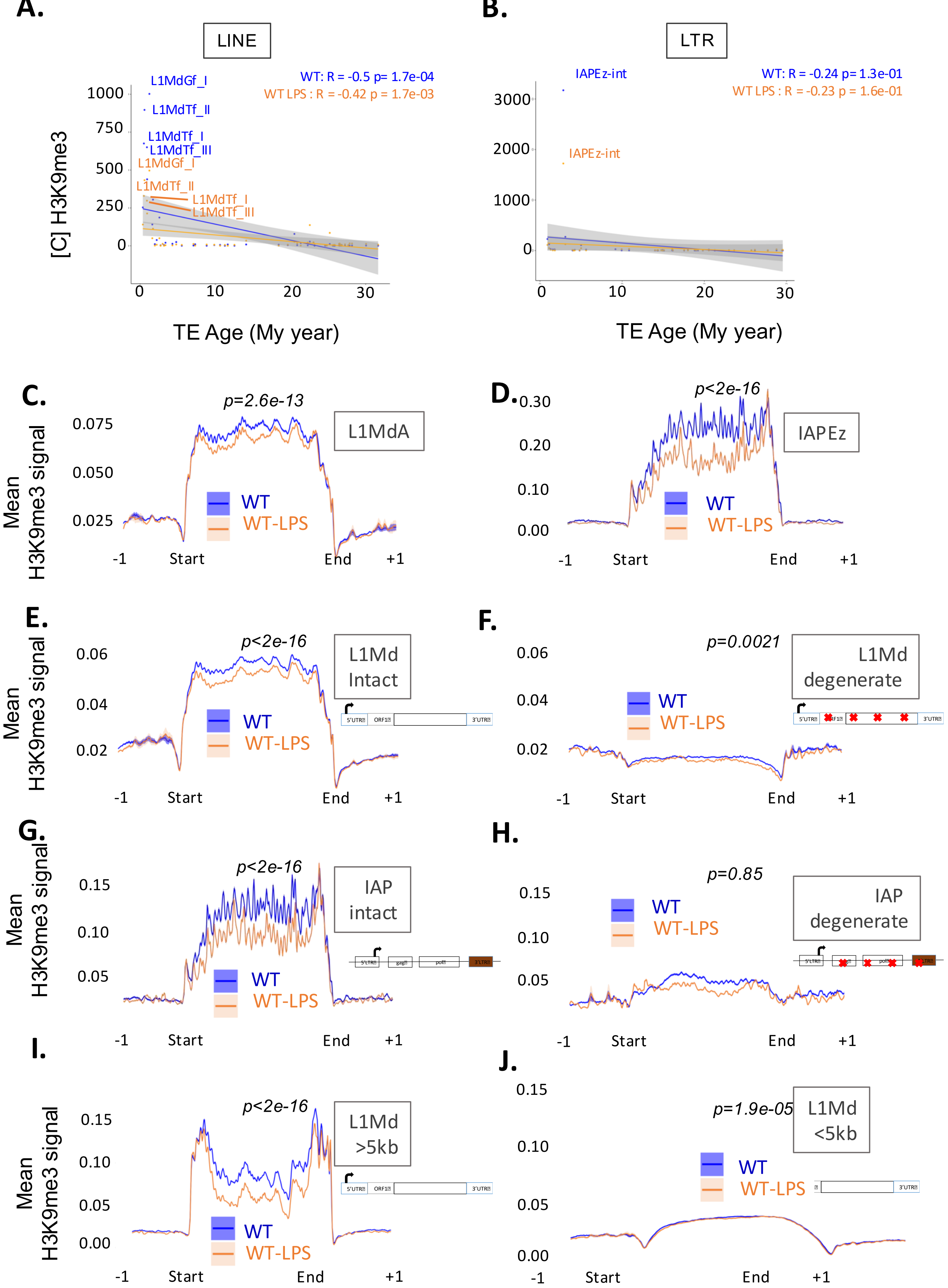


**Fig 2. H3K9me3 loss mostly occurs at LINE and LTR, and especially at L1Md and IAP, the most recent subfamilies**

**Figure S2 (related to figure 2). (A-B)** Correlation plot representing H3K9me3 concentration quantified at differentially enriched LINE (A) or LTR (B) elements retrieved in Figure 2A *vs* their age in million years (My) in WT and WT-LPS conditions. R, Pearson correlation coefficient; p, pvalue. **(C-J)** Plot profiles of the mean +/- SEM H3K9me3 enrichment along L1MdA (C) and IAPEz (D) sequences, intact (E) vs degenerate (F) L1Md, intact (G) vs degenerate (H) IAP, (intact meaning >97% conservation as previously defined (Bulut-Karslioglu et al., 2014)), and >5kb (I) and <5kb (J) L1Md sequences +/- 1kb flanking regions in WT-LPS (orange) *vs* WT (blue) conditions. (2-3 samples per condition). Red crosses symbolize sequence variation towards concensus sequence in degenerate L1Md or IAP.

FIGURE S3

Related to figure 3

Stolz et al.

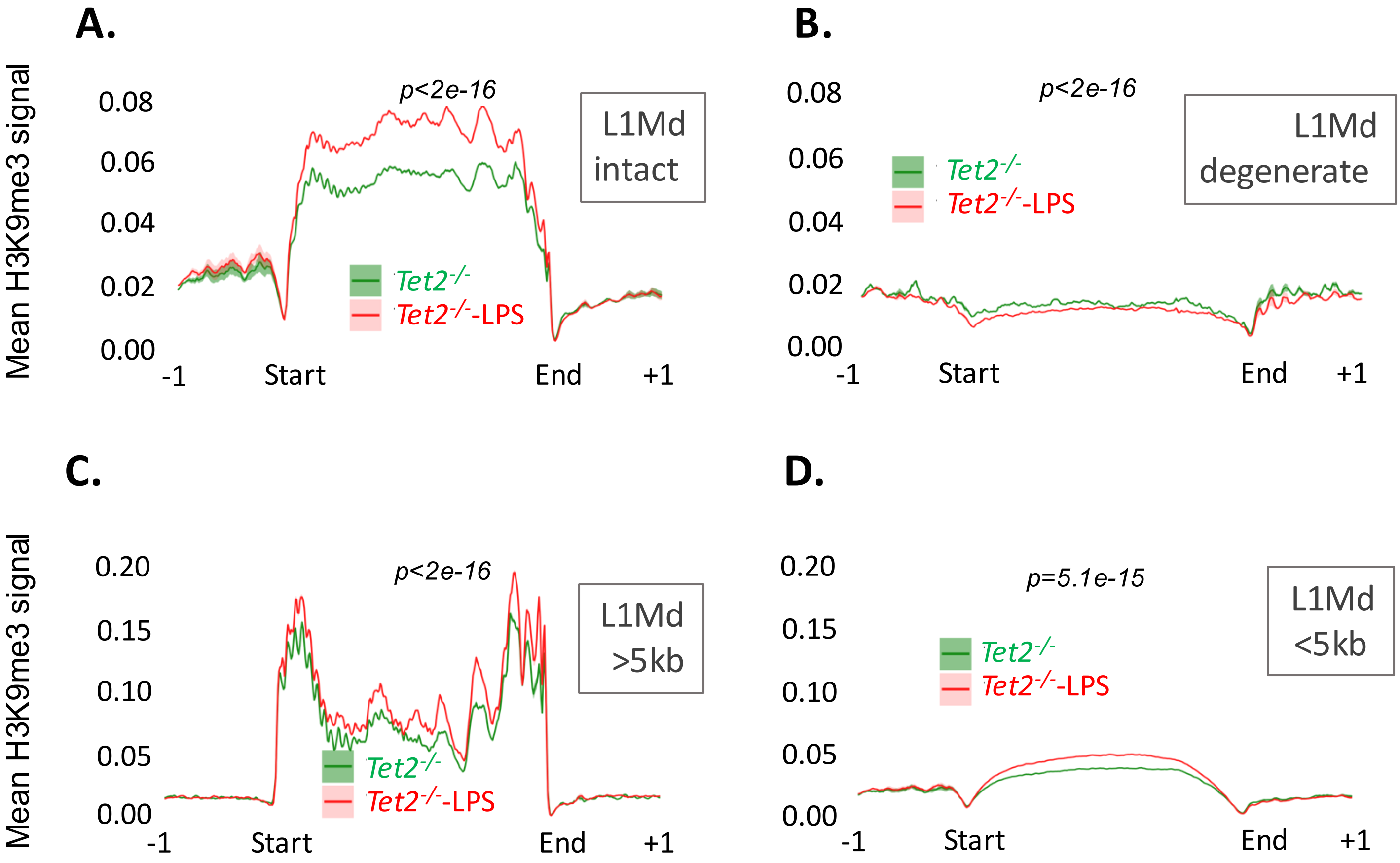


**Figure S3 (related to figure 3).** **(A-D)** Plot profile of the mean +/- SEM H3K9me3 enrichment along intact (A) or degenerate (B) and >5kb (C) or <5kb (D) L1Md sequences +/- 1kb flanking regions in *Tet2*$^{-/-}$-LPS (red) and *Tet2*$^{-/-}$ (green) conditions (2 samples per condition).

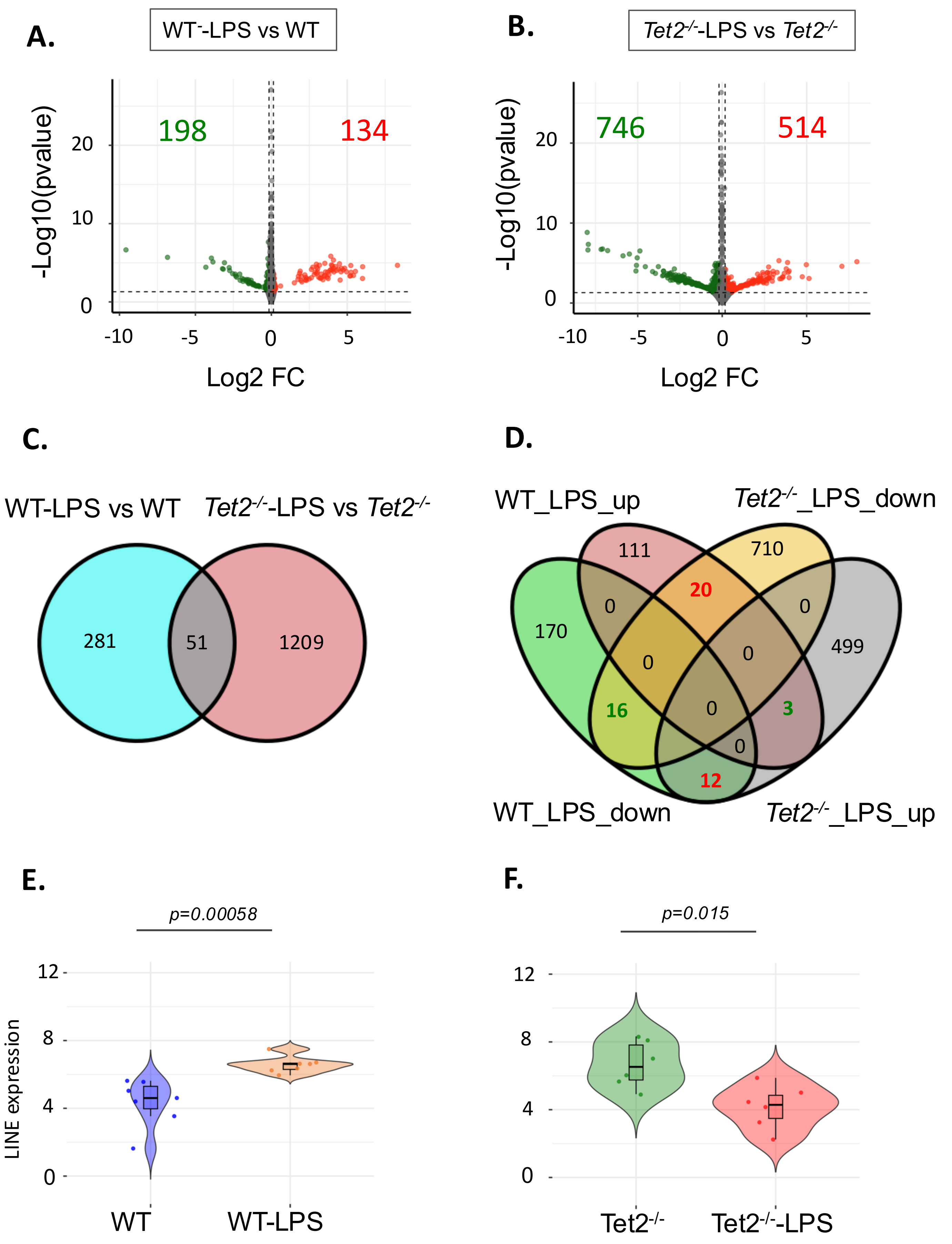
A.
WT⁻-LPS vs WT
198
134
-Log10(pvalue)
Log2 FC
B.
Tet2-/--LPS vs Tet2-/-
746
514
-Log10(pvalue)
Log2 FC
C.
WT-LPS vs WT
Tet2-/--LPS vs Tet2-/-
281
51
1209
D.
WT_LPS_up
Tet2-/-_LPS_down
WT_LPS_down
Tet2-/-_LPS_up
111
710
20
170
499
16
3
12
E.
p=0.00058
LINE expression
WT
WT-LPS
F.
p=0.015
Tet2-/-
Tet2-/--LPS

**Figure S4 (related to Figure 4): (A-B)** Volcano plot of gene expression changes between WT-LPS and WT (A) and between *Tet2*$^{-/-}$-LPS and *Tet2*$^{-/-}$ (B) conditions. The vertical axis represents the –log10(pValue) and the horizontal axis the log2 fold change (FC). Genes presenting a significant (*p<0.05, 0.9<FC<1.1)* increase (red) or decrease (green) expression are shown. **(C-D)** Venn diagram of DEGs (*p<0.05, 0.9<FC<1.1*) in observed between WT-LPS vs WT and *Tet2*$^{-/-}$-LPS vs *Tet2*$^{-/-}$comparisons. In (C), all DEGs are shown, regardless of the direction of change and in (D), the genes have been separated according to the direction of change observed upon LPS treatment, with colours indicating the number of genes inversely (red) or commonly (green) deregulated in WT and *Tet2*$^{-/-}$ HSCs. **(E-F)** violin plot representing L1 subfamilies expression for each L1 subfamily significantly (p<0.05) differentially expressed in WT-LPS vs WT (E) and *Tet2*$^{-/-}$-LPS vs *Tet2*$^{-/-}$ (F) conditions.

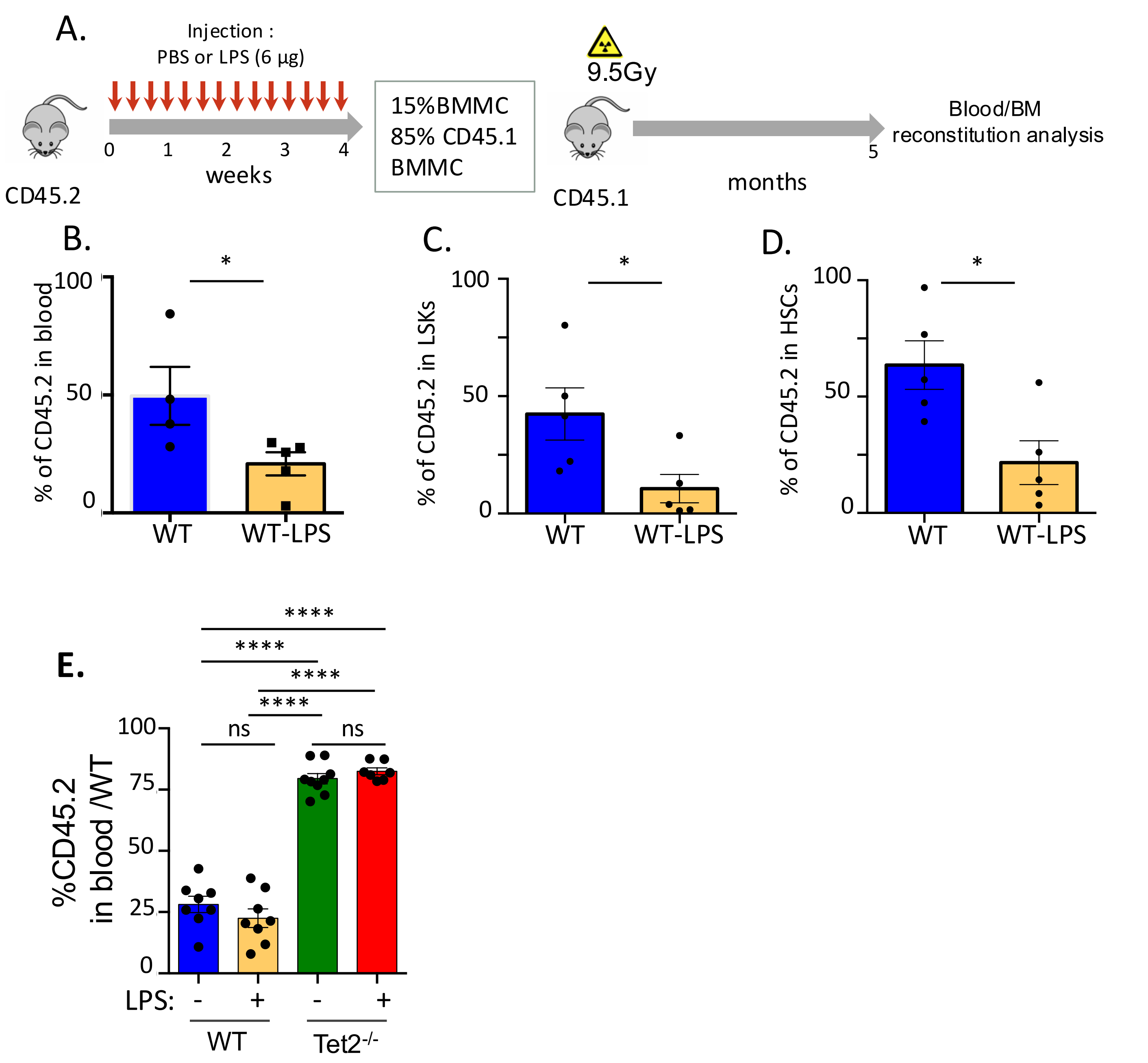


**Figure S5 (related to Figure 6). (A)** Experimental design for reconstitution experiments using BM from mice treated with PBS (WT) or LPS (WT-LPS) every two days for 30 days; BMMC, bone marrow mononuclear cells. **(B)** Percentage of CD45.2 donor contribution in blood in mice transplanted with WT or WT-LPS cells 5 months after reconstitution. **(C-D)** Percentage of LSK CD45.2+ (C) or LSK CD34- Flk2- CD45.2+ (HSC) (D) donor contribution in BM 5 months after reconstitution. 5 mice per group. Mann Whitney test $^*p<0.05$. **(E)** Percentage of CD45.2 cells in blood after LPS treatment in mice engrafted with 10% WT or *Tet2*$^{-/-}$ BM cells 3 months after the end of the treatment. 4-5 mice per group in 2 independent experiments. One-way ANOVA with Tukey's multiple comparison test.

FIGURE S6 Related to figure 7 Stolz et al.

**A.**

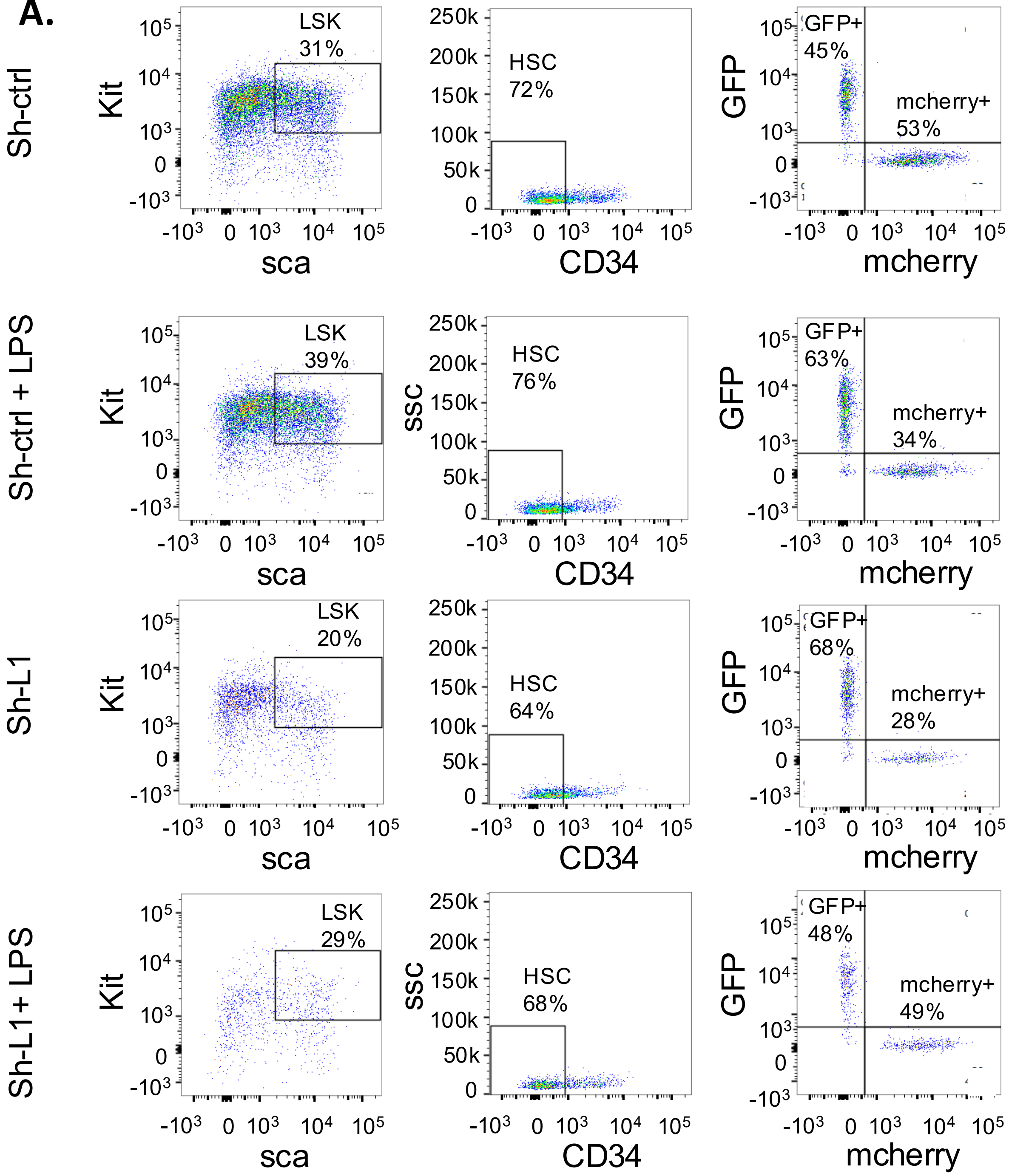

**B.**

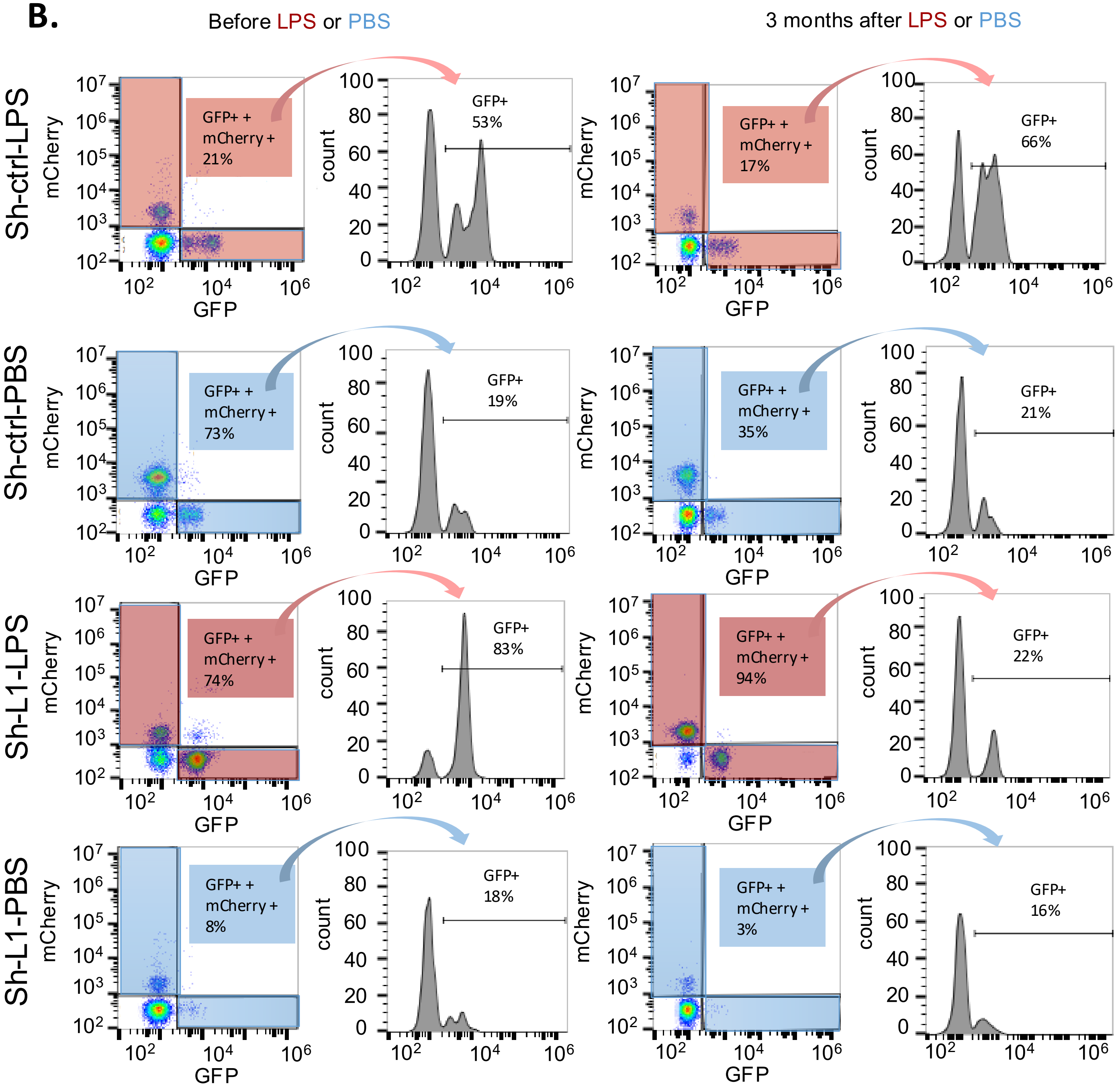


**(A)** Gating strategy for studying the competition between *Tet2*$^{-/-}$ cells transduced with an sh-control-GFP and WT HSCs (CD34- low) transduced with an sh-control-mCherry or sh-LINE-mcherry in the presence or absence of LPS during 3 days in liquid culture in vitro. **(B)** Gating strategy for studying the competition between *Tet2*$^{-/-}$ cells transduced with an sh-control-GFP and WT cells transduced with an sh-control-mCherry or sh-LINE-mcherry in the peripheral blood of CD45.2 recipient mice before (T-1) or 3 months after LPS or PBS treatment.
